# Semirandom DNA adducts regulate a filamentous defence-associated reverse transcriptase

**DOI:** 10.64898/2026.03.07.710306

**Authors:** Nolan Neville, Nicole V. Johnson, Edwin E. Escobar, Chang-Hwa Chiang, Albana Nreca, Sean R. Johnson, Nan Dai, Andy Hanneman, Ivan R. Corrêa, Jason S. McLellan, Robert J. Trachman

## Abstract

Retrons and several defence-associated reverse transcriptases (DRTs) synthesize non-genomic DNA for bacteriophage immunity. In some instances, this non-genomic DNA is of undefined, semirandom sequence. How undefined DNA sequences impart antiphage defence is not known. Here we report the cryo-EM structure and functional characterization of the DRT1 antiphage defence system. We show that DRT1 performs template-free, protein-primed DNA synthesis to generate semirandom DNA adducts. DNA synthesis activates the nitrilase domain of DRT1 while DNA adducts drive assembly of quiescent DRT1 filaments. Filamentous DRT1 is comprised of domain-swapped C-termini that are entwined, forming pseudoknots between tetrameric stacks. This configuration occludes the apo-nitrilase active site, resulting in a dormant state. Bacteriophage escape mutants identify a T4 single-stranded DNA helicase required for DRT1 activity. Functionally, DRT1 resembles a minimal retron where a single gene produces an RT, effector, and non-genomic antitoxin DNA.

## Introduction

Bacteria have evolved diverse pathways to contest bacteriophage. One broadly implemented defence strategy is to degrade foreign nucleic acids^1-3^. Contrasting with this tactic, defence-associated reverse transcriptases (DRT) synthesize DNA to impart immunity. DRTs are part of the so-called unknown group (UG) of RTs, which are closely related to the Abortive infection (Abi) RTs. Collectively, these are referred to as UG/Abi RT systems. Groundbreaking work by several research teams^4-6^ identified dozens of UG/Abi RT families that are categorized into three major classes based on functional gene annotation or domain association^6^. Class 1 UG/Abi RTs homo-oligomerize to perform template-free, protein-primed DNA synthesis^7,8^. This process results in a homo-oligomeric protein core with DNA adducts of seemingly random sequences. The function of undefined single-stranded DNA (ssDNA) in antiphage defence has yet to be determined.

Like retrons^9-11^, Class 2 UG/Abi RT operons encode an RT along with a noncoding RNA and sometimes an additional protein. Despite this similarity, the two characterized Class 2 DRTs, DRT2 and DRT9, are mechanistically distinct and unparalleled in nature. Retron RTs produce multicopy single-stranded DNA (msDNA) as an autoregulatory antitoxin that stabilizes filaments and senses phage infection^12-14^, while Class 2 DRTs produce DNA templates encoding toxic products^15-18^.

Despite wide distribution across diverse bacterial taxa, Class 3 UG/Abi RTs are the least characterized. Class 3 UG/Abi RTs are defined by multidomain proteins bearing an RT domain and either a nitrilase or phosphohydrolase domain^6^. These systems are intriguing as neither nitrilases nor phosphohydrolases are common to antiphage defence systems^5,19^. These attributes suggest that Class 3 UG/Abi RTs function through an unprecedented mechanism.

Using cryogenic electron microscopy (cryo-EM) and functional assays we investigate the mechanism of the Class 3 DRT1 antiphage defence system. We demonstrate that Class 3 DRTs perform protein-primed, template-free DNA synthesis to impart immunity to bacteriophage. DRT1-DNA adducts serve to activate the nitrilase domain by inducing a tetrameric state while simultaneously imparting a dormant filamentous state. Bacteriophage escape mutants identified the T4 5’→3’ single-stranded DNA helicase, Dda, as part of the activating network of DRT1. Collectively, our results reveal how DRT1-DNA adducts promote a pre-active state primed for inducing programmed cell death upon infection.

## Results

### A diverse class of DRTs perform template-free protein-primed DNA synthesis

Defence-associated reverse transcriptases (DRTs) within the UG/Abi RTs are grouped into three classes defined by their association with conserved domains^6^. Phylogenetic analysis of 31 previously annotated UG/Abi RT families and retrons revealed that Class 3 DRTs reside in a distinct clade (**Fig. 1a**), suggesting a unique structure and mechanism. Indeed, the predicted structure of the Class 3 DRT1 by AlphaFold2^20^ (pLDDT = 86.2) is distinct from those of Class 1 and Class 2 DRTs with the RT and nitrilase domains forming a compact globular shape (**Fig. 1a**). These data support a structural role for these domains but do not address any functional enzymatic requirement for either the RT or nitrilase domain in antiphage defence.

**Fig. 1.**
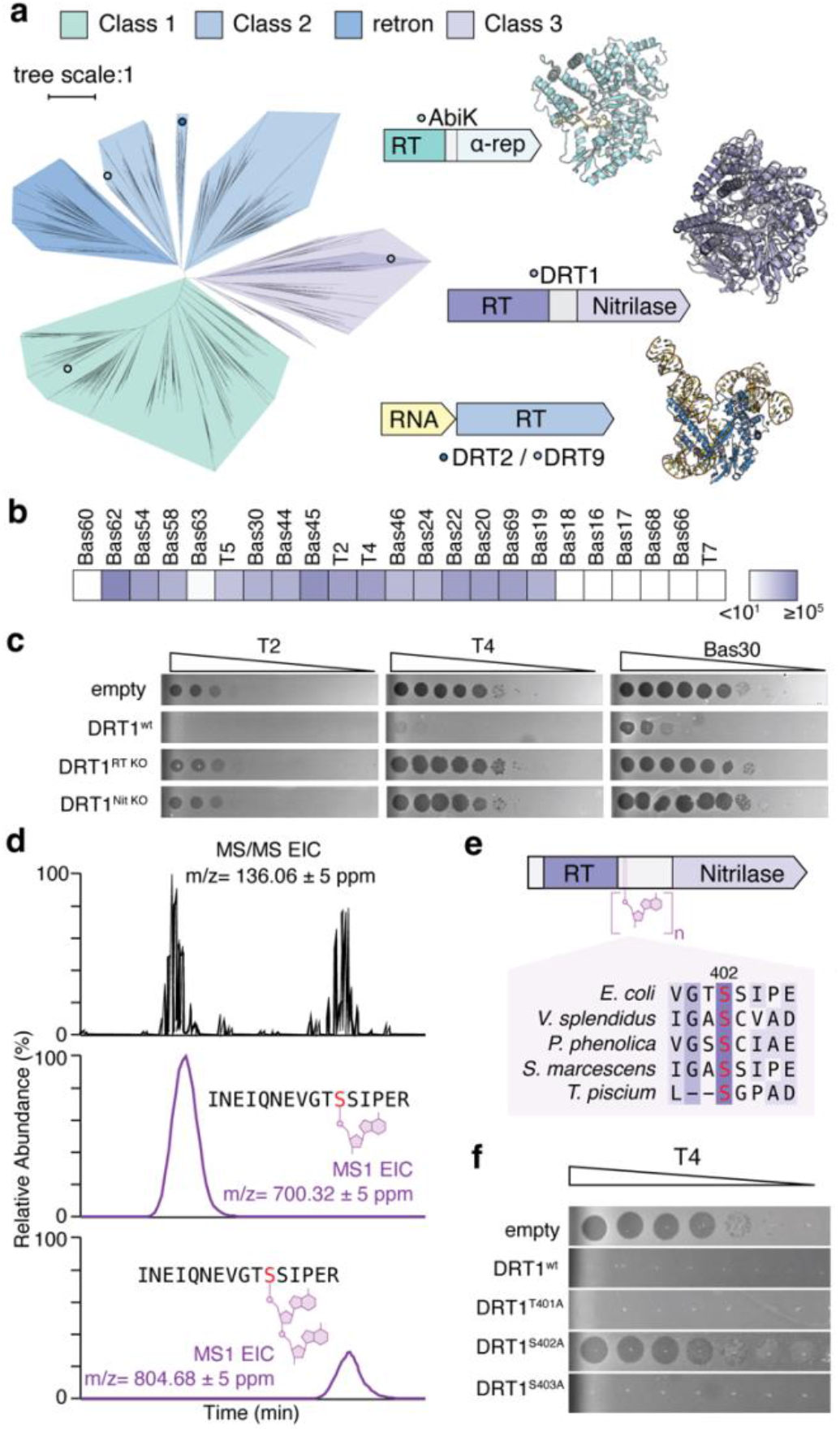
Antiphage defence via protein-primed DNA synthesis. **a)** Phylogenetic analysis of 31 UG/Abi RT families and retrons^4^. Clades are coloured by UG/Abi RT Class as described by Mestre et al. ^6^. Cartoon representation of enzymes and enzyme complexes represents published structures of AbiK ^7^ and DRT2^15^ or the AlphaFold2^20^ model for DRT1. **b)** Heat map representing plaquing of tenfold serial dilutions of bacteriophage on lawns of *E. coli* MG1655. Darker shades represent greater protection against bacteriophage. **c)** Plaquing of tenfold bacteriophage serial dilutions on lawns of *E. coli* MG1655 with control (empty vector), DRT1^wt^, and active site mutants. **d)** LC-MS traces of trypsin-digested DRT1 with detection of adenine diagnostic fragment. **e)** Cartoon representation of DRT1 identifying relative location of serine priming site. Limited sequence alignment window focusing on conserved serine 402. **f)** Plaquing of tenfold bacteriophage serial dilutions on lawns of *E. coli* MG1655 with control (empty vector) and expressed DRT1 and DRT1 priming-site mutants.

We performed spot titer assays to test the function of DRT1 and its individual domains using a bacteriophage library composed of T-phages and a subset of the Basel collection^21^. In the surrogate host *E. coli* MG1655, DRT1 provides robust antiphage defence against a diverse subset of representative bacteriophage (**Fig. 1b**). Mutagenesis of either the DRT1 RT active site residues D324N/D325N (DRT1^RTKO^) or nitrilase active site residue C1116A (DRT1^NitKO^) abolished immunity to bacteriophage (**Fig. 1c**). Consistent with abortive infection, DRT1 only provided immunity at low multiplicities of infection (MOI) and reduced bacteriophage burst size while maintaining colony survival rate (**Extended Data Fig. 1a-c**). Like the RT-toprim fusion of retron-Eco2 ^22^, enzymatic activity of the RT and effector domains of DRT1 is required to induce abortive infection. Together, these data support a defence mechanism distinct from either Class 1 or Class 2 DRTs.

To experimentally test the biochemical properties of Class 3 DRTs we expressed and purified DRT1, DRT5S and DRT5L. Reminiscent of the Class 1 UG/Abi RTs, Class 3 DRTs synthesized DNA upon incubation with dNTPs in the absence of a supplemented RNA template and in the presence of RNase A (**Extended Data Fig. 1d**). This feature is indicative of protein-primed DNA synthesis, a mechanism observed in bacteriophage polymerases^23,24^ and antiphage defence systems^7,8,18,25^, which results in covalent fusion between protein and DNA. Therefore, we chose to test if DNA is covalently bound to DRT1 and DRT5. Post reaction with dNTPs, DNA products were only resolved by electrophoresis for samples digested with proteinase K (**Extended Data Fig. 1e**). Furthermore, both DRT1^wt^ and DRT1^NitKO^ synthesized DNA **(Extended Data Fig. 1f)**. Additionally, intact mass spectrometry of DRT1^wt^ resolved masses of apo-DRT1 and DRT1• dAMP (**Extended Data Fig. 3,4**). These data support that Class 3 DRTs produce covalent DNA adducts through template-free, protein-primed, DNA synthesis independent of nitrilase activity.

Focusing on DRT1, we sought to determine the amino acid priming site by developing a tandem mass spectrometry method following a similar approach developed for detection of O-glycosylation in proteins^26^. Following tryptic digestion, an HCD-triggered-EThcD MS/MS method was used to observe nucleotide-specific diagnostic fragment ions confirming covalently bound nucleotides. This analysis revealed S402 as the priming site, as evidenced by the detection of four key fragment ions on a single dA-modified peptide. Up to four deoxyadenosine residues were observed on this priming site (**Fig. 1d; Extended Data Fig. 4; Extended Data Fig. 3**). S402 is highly conserved within the DRT1 family (**Fig. 1e**), but not in other Class 3 DRTs. To validate the priming site, we mutated S402 and the adjacent residues T401 and S403 to alanine. The S402A mutation completely abolished bacteriophage immunity, while T401A and S403A mutations had no effect (**Fig. 1f**). The use of a single serine residue for DRT1 priming is a notable distinction from other protein-primed UG/Abi RTs.

### DNA adducts promote a dormant filamentous state of DRT1

DRT1 exhibited unusual properties during purification. Size exclusion chromatography (SEC) revealed a broad elution peak corresponding to a mass range of ∼440 kDa to ∼3 MDa with a second peak eluting with an estimated mass of ∼6 MDa (**Extended Data Fig. 2a**). Analysis of individual SEC fractions by SDS-PAGE under reducing conditions revealed only the presence of DRT1^wt^ (**Extended Data Fig. 2b**). We performed mass photometry on DRT1^wt^, DRT1^NitKO^, and DRT1^RTKO^ to further understand the mass distribution of DRT1. The mass profile of natively prepared DRT1^wt^ and DRT1^NitKO^ comprised eight or more species with average masses approximating to multiples of tetrameric DRT1 (**Fig. 2c**). Upon reaction with dNTPs, the mass profiles became monodispersed. As confirmed by SDS-PAGE (**Extended Data Fig. 2b**), these data are consistent with protein-primed DNA synthesis causing shifts to high molecular weight. The DRT1^RTKO^ mutant was monomeric (**Fig. 2c**), confirming the requirement of DRT1-DNA adducts to form high-order oligomeric states. The reaction of DRT1^wt^ with dideoxy ATP did not alter the oligomer distribution (**Extended Data Fig. 2c**). Together, these data indicate that DNA adducts are required for filamentation.

**Fig. 2.**
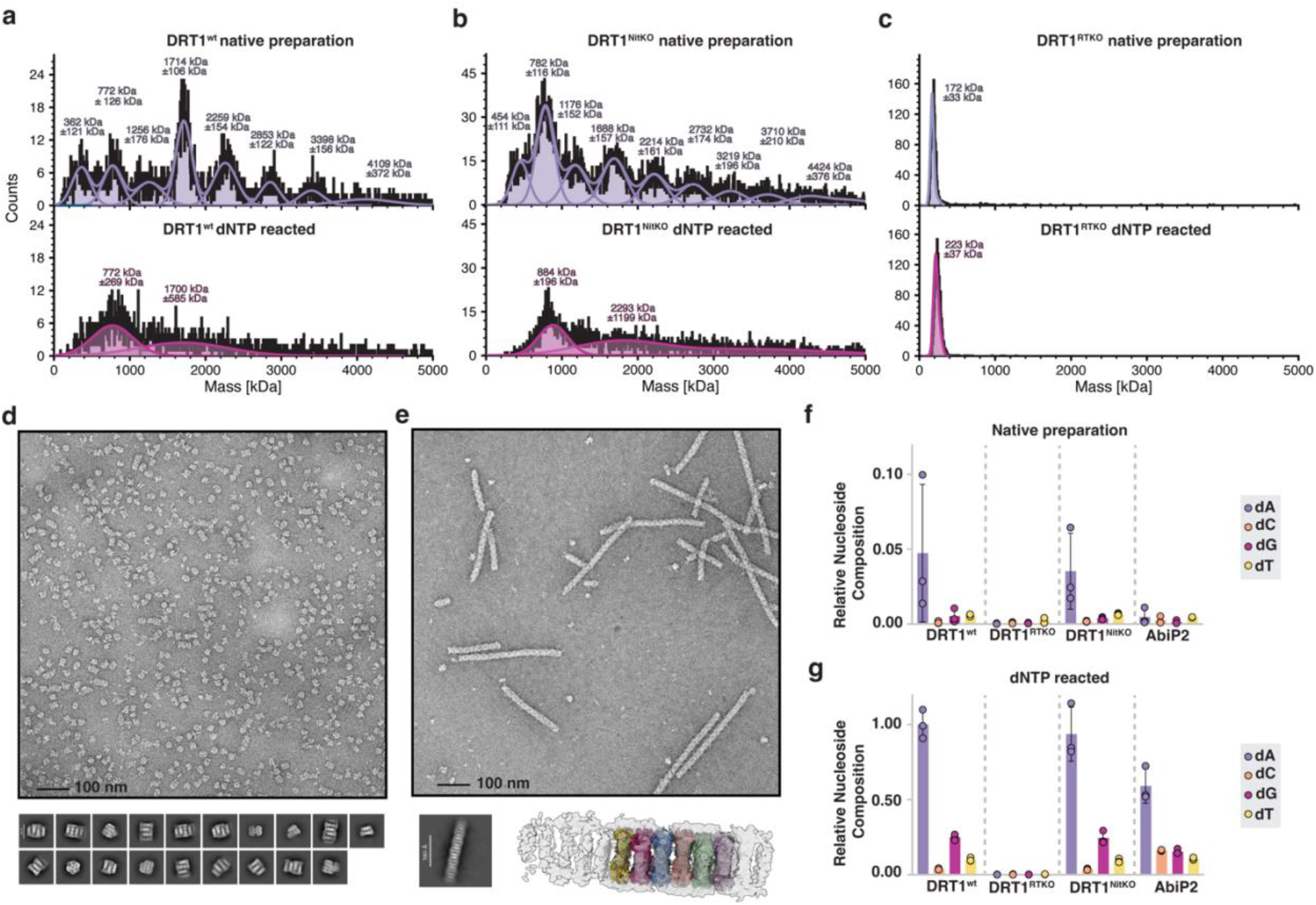
DNA adducts drive filament formation. **a)** Mass photometry of DRT1^wt^, **b)** DRT1^NitKO^, **c)** DRT1^RTKO^ from native preparation (purple) and dNTP reacted (magenta) samples. Fits to read counts are shown in solid line with average and standard error of masses labeled above peaks. **d)** Negative stain electron micrograph of DRT1 from native preparation. 2-D class averages are shown below micrograph image. **e)** Negative stain electron micrograph of dNTP-reacted DRT1. 2D-class average of filaments is shown below micrograph along with 3D reconstruction of filament with AlphaFold2-predicted DRT1 tetramers (coloured models) fit to electron density. **f)** Relative nucleoside composition of DRT1^wt^, DRT1 variants, and AbiP2 from native preparation. **g)** Relative nucleoside composition of dNTP-reacted DRT1^wt^, DRT1 variants, and AbiP2.

Next, we performed negative stain electron microscopy on unreacted and dNTP-reacted DRT1 to understand the organization of DRT1 oligomers and how DNA synthesis alters these assemblies. Electron micrographs of natively prepared DRT1 revealed a heterogenous population of protein assemblies (**Fig. 2d**). The resulting nineteen 2D population class-averages show a distribution of stacked planar assemblies forming truncated filaments tens of nanometers in length. The reaction of DRT1 with dNTPs increased the filament length to >100 nm (**Fig. 2e**). Consistent with the tetrameric repeats observed by mass photometry, AlphaFold2 predicted tetramers of DRT1 fit to the 3D reconstruction of the reacted sample. Furthermore, nucleobase composition of the DNA adducts did not impact filament formation (**Extended Data Fig. 5a-c**) despite DRT1 exhibiting a higher propensity to incorporate dATP in cells and *in vitro* (**Fig. 2f,g; Extended Data Fig. 5d**). These data demonstrate that DRT1 expressed in *E. coli* contains DNA adducts and forms filamentous structures prior to bacteriophage infection. Given the expression of DRT1^wt^, or the active-site mutants, does not alter growth kinetics of *E. coli* (**Extended Data Fig. 5e**) the filamentous and monomeric forms of DRT1 are not toxic, requiring phage infection to induce cell death.

### Overall structure of DRT1

To further investigate the molecular basis for DRT1 filament assembly, we performed single-particle cryo-EM on dNTP-reacted DRT1^wt^. Ab initio reconstruction and heterogeneous refinement of particles extracted from the filaments yielded a high-quality 3D reconstruction revealing a right-handed helical structure composed of stacked DRT1 tetramers. A helical symmetry search performed in cryoSPARC yielded a high-confidence candidate symmetry pair (twist = -121.22°, rise = 59.13Å) corresponding to a repeating octameric DRT1 unit. These parameters were used in subsequent helical refinement. A final reconstruction of 140,634 particles with additionally imposed D2 point group symmetry yielded a 2.6 Å resolution reconstruction of the DRT1 filament **(Fig 3a; Extended Data Table 2; Extended Data Fig. 6)**. The high-resolution cryo-EM reconstruction allowed for unambiguous model building of the DRT1 octamer. Residues 382-478, including the priming site residue S402, and 837-846 were not resolved in the map.

The DRT1 tetramer is defined with monomeric units arranged clockwise as α, β, γ, and δ with protomer interfaces ϕ and χ **(Fig. 3b)**. Tetramers assemble with the RT and alpha-helical repeat domains forming the filament periphery and the nitrilase domains forming the core. Intra-tetramer contacts are exclusively mediated by residues from the nitrilase domain. In total <u>4,278</u> Å^2^ of surface area is buried between the nitrilase ϕ (<u>2,614</u> Å^2^) and χ (<u>1,664</u> Å^2^) faces. The octameric subunit of DRT1 filaments is composed of tetramers stacked along a central axis. Successive tetramers are flipped by 180◦ along the plane of the tetramer and rotated by 60.61◦ (**Fig. 3b**). This repeating arrangement results in inter-tetramer Van der Waals and electrostatic interactions between RT and alpha-helical repeat domain between residues 689-692 and 696-715 to 199-204, 332-336, and 431-437.

**Fig. 3.**
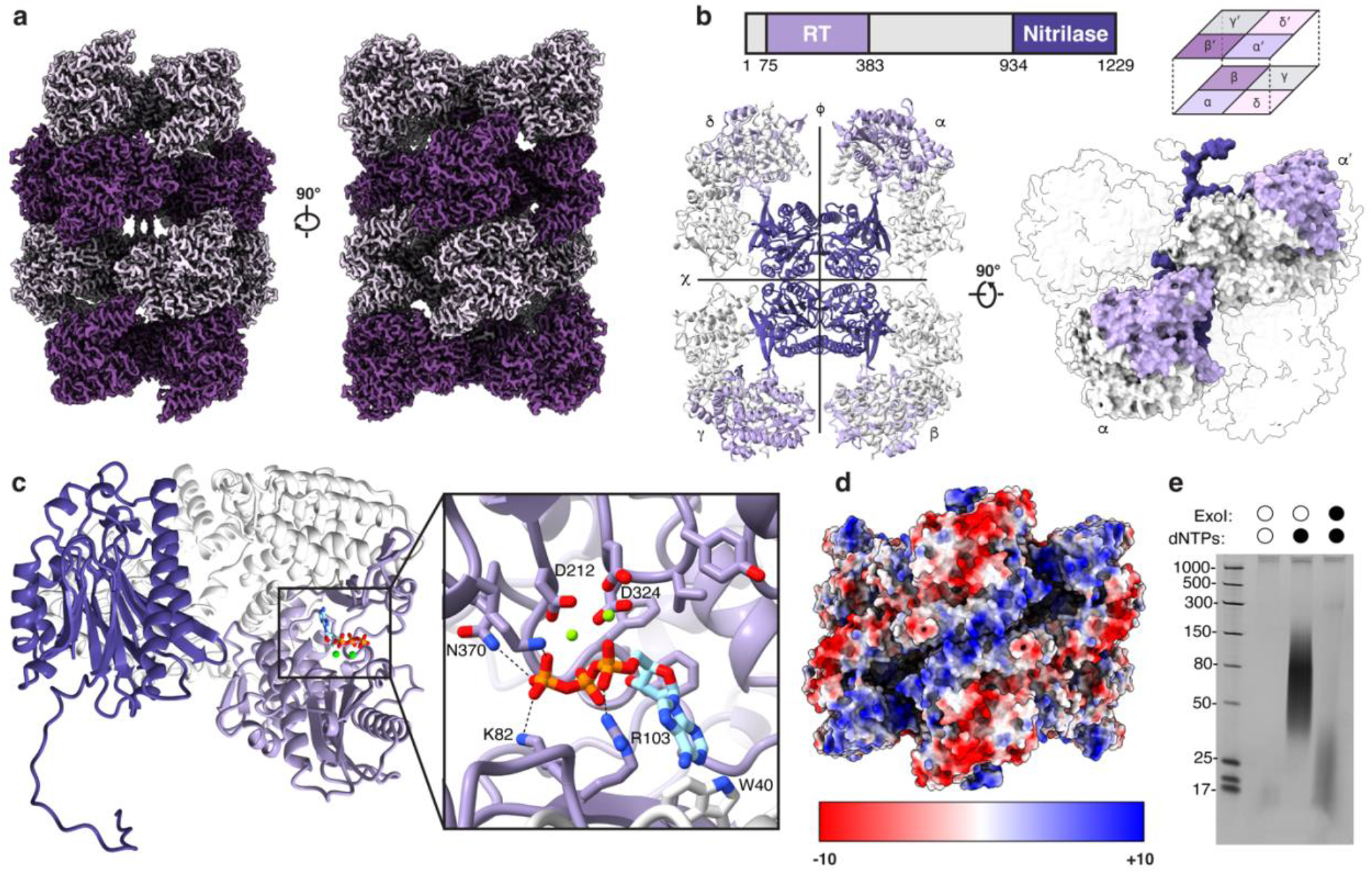
Cryo-EM structure of the DRT1 filament. **a)** Electron density map of the DRT1 filament. **b)** Arrangement and nomenclature of DRT1 protomers. Protomers are labeled α−δ in the tetramer model. Surface representation showing inter-protomer interactions between tetramer stacks is shown on right. **c)** Cartoon representation of DRT1 monomer coloured by domain with inset showing dATP substrate in RT active site. **d)** Electrostatic surface of the DRT1 octamer. **e)** Urea-PAGE of DRT1 DNA adducts from native preparation, post-reaction with dNTPs, and post-reaction with dNTPs and Exonuclease I digest.

Despite our determination that ssDNA synthesis is responsible for stabilization of the filamentous structure, we were unable to resolve map density corresponding to ssDNA on the DRT1 surface. The only nucleic acid resolved in our structure is a single deoxyadenosine triphosphate molecule coordinated by two Mg^2+^ ions in the RT active site **(Fig. 3c)**. DNA adduct interactions are likely electrostatic given the lack of sequence dependence on filamentation ^27^. To further investigate how ssDNA may interact with the DRT1 filament the electrostatic potential was calculated for the DRT1 octamer. The electrostatic surface potential revealed an uninterrupted helical patch of positive potential at the interface between tetramers with an approximate length of 130 Å per tetramer **(Fig. 3d)**. Natively purified or Exonuclease I treated DRT1 results in DNA adducts of ∼17-30 nucleotides **(Fig. 3e)**. These data are consistent with DNA adduct nuclease protection correlating in length with the basic helical patch.

### C-terminal domain swapping

In addition to extensive inter-tetramer contacts, the final nineteen C-terminal (K1211-H1229) residues extend across the interstitial space of the octamer resulting in two domain-swapped tetramers per octamer **(Fig. 4a)**. The C-termini from protomers on opposite tetramer stacks wrap around each other in an antiparallel orientation to form a pseudoknot (**Fig. 4a**,**b**). The inter C-terminal domain contact is stabilized by cation-π interactions between R1210 and Y1215 and van der Waals interactions between I1212 and Y1215 **(Fig. 4a)**. This conformation positions C-terminal residues 1218-1221 over the apo-nitrilase binding pocket of the protomer adjacent to pseudoknot pair (**Fig. 4c**). This produces a closed-lid conformation similar to those observed between adjacent protomers in bacterial and plant nitrilases **(Fig. 4d)**^28,29^ with only modest differences in the conformation of the catalytic tetrad (**Fig. 4e**).

**Fig. 4.**
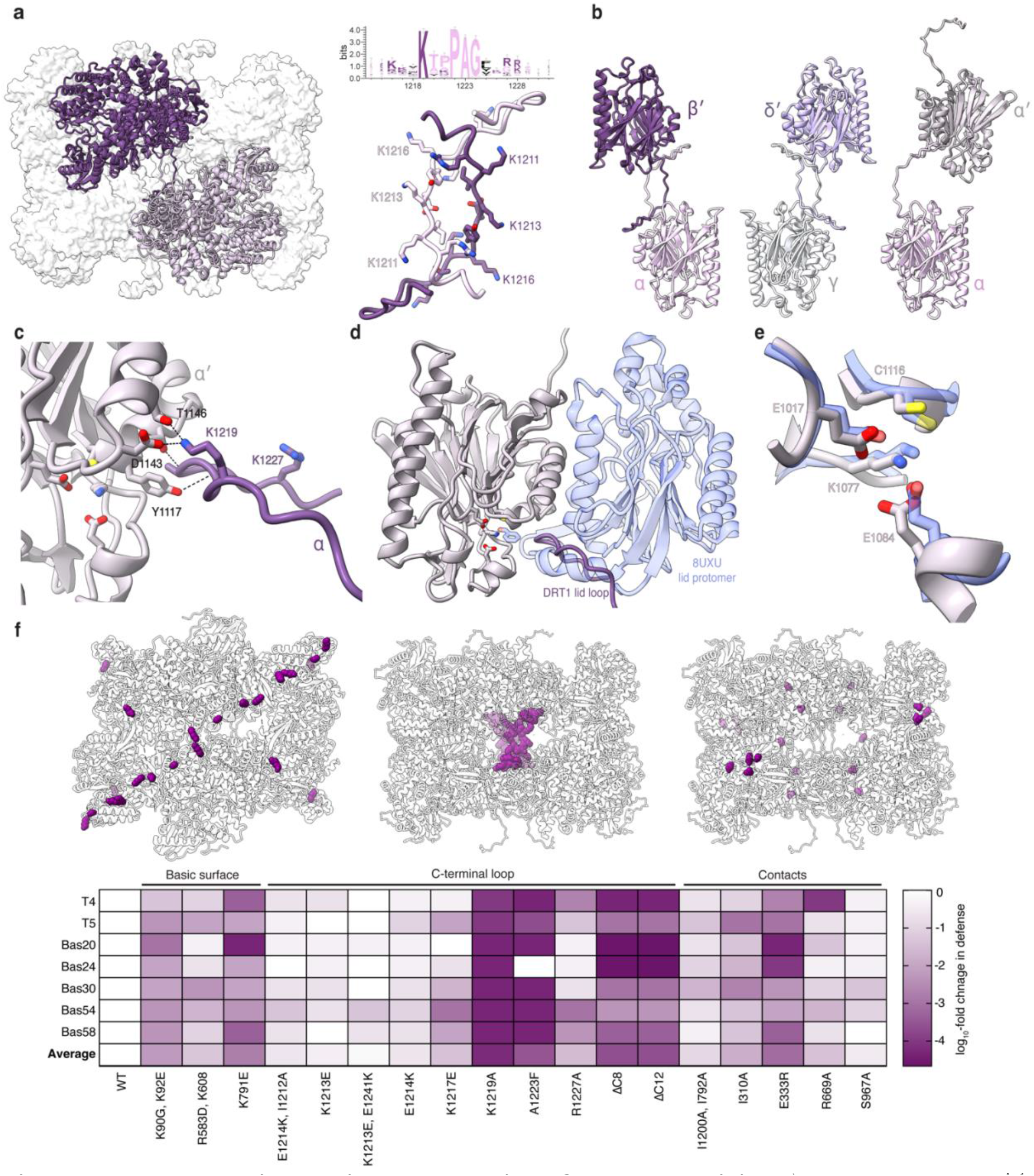
Inter-tetramer interactions are required for DRT1 activity. **a**) DRT1 octamer with domain-swapped C-termini shown in inset panel. Sequence logo of the C-terminal residues generated from alignment of 174 DRT1 sequences. **b**) Orientation of protomers within with domain swapping interactions. A prime (’) symbol indicates a protomer from a different tetramer stack. **c)** Interactions between proximal residues to the nitrilase active site (light pink) and swapped C-terminus of adjacent protomer (purple). **d)** Closed lid conformation of DRT1 nitrilase domain (light pink) superimposed with the dimeric unit of *Rhodococcus* sp. V51B (PDB: 8UXU) and its benzaldehyde substrate (blue). The domain swapped C-terminus of a DRT1 protomer is shown in purple, and the nitrilase active site catalytic tetrad is shown as sticks. **e**) Superposition of the DRT1 nitrilase active site catalytic tetrad (light pink) with that of a nitrilase filament from *Rhodococcus* sp. V51B (PDB: 8UXU) (blue). **f**) Structure-guided mutations suggested to disrupt DRT1 activity, with mutation locations shown as purple spheres in the structures with heatmap of phage plaquing assay relative to DRT1^wt^. Darker shades indicate loss of immunity.

To test the structural and functional significance of DRT1 filament features, we performed a spot-titer assay on mutants of DRT1^wt^, evaluating plaquing efficiency of *E. coli* inoculated with one of seven phages. Single or double amino acid substitutions were selected to target basic surface residues hypothesized to interact with DNA adducts, residues involved in C-terminal domain swapping, and inter-tetramer contacts **(Fig. 4d)**. The most striking reduction in antiphage defence resulted from substitutions to conserved residues in the C-terminal loop region, with reduction in plaque formation up to four orders of magnitude higher than DRT1^wt^. Point substitutions K1219A and A1223F in this region reduced plaquing to levels comparable to C-terminal deletions, Δ1217-1229 (ΔC12) and Δ1221-1229 (ΔC8), revealing these residues to be essential for DRT1 activity. By contrast, non-conserved residues in the C-terminal loop resulted in less than or equal to a 10-fold reduction in plaquing. Except for E333R, substitutions targeting tetrameric contacts had modest impact on plaquing. Maintenance of antiphage defence activity in these mutants is unsurprising due to the substantial amount of buried surface area between DRT1 tetramers.

### Viral determinants of DRT1 immunity

To understand bacteriophage gene association and response to DRT1, we performed RNA-seq on T4- and T5-infected *E. coli* MG1655 expressing DRT1. Ten T4 mid-stage genes with known transcriptional or replication functions were significantly upregulated in DRT1 expressing strains, while five genes encoding structural proteins and nucleotide metabolism were downregulated (**Extended Data Fig. 7a**). Conversely, bacteriophage T5 only showed significant downregulation of genes encoding structural proteins, nucleotide metabolism, and endolysin (**Extended Data Fig. 7b**). Noteworthy changes to transcription profiles are the down-regulated genes nrdC.8 of T4 and nrdA of T5. Such nucleotide reductases have been shown to stimulate DRT9^17^. Both RT and nitrilase catalytic activity are required for significant alteration to relative bacteriophage transcriptome profiles (**Extended Data Fig. 7)**. Together, these data suggest DRT1 activation occurs during mid-stage of bacteriophage infection.

DRT9 produces long DNA adducts upon bacteriophage infection. Given mRNA levels of transcriptional and replication enzymes are upregulated during bacteriophage T4 infection we postulated that DRT1 may function by a similar mechanism to DRT9. To determine whether phage infection alters the length or composition of the DNA adducts, we applied a variation of cDNA immunoprecipitation sequencing (cDIP-seq)^16,18^. First, we validated that adding either MBP or 3xFLAG tags to DRT1 did not affect antiphage defence (**Extended Data Fig. 8a**). Since nucleoside analysis indicated that DRT1 ssDNA is comprised of at least 70% A (**Fig. 2f,g**), we extracted and analyzed unmapped cDIP-seq reads containing ≥70% adenosine (A-rich reads) obtained from DRT1 immunoprecipitation in the presence or absence of T4 bacteriophage infection. No increase in A-rich ssDNA or overall unmapped reads was observed in DRT1-expressing cultures upon T4 infection (**Fig. 5a; Extended Data Fig. 8b**). Indeed, only the positive control consisting of purified untagged DRT1 reacted *in vitro* with dNTPs showed a significant enrichment of A-rich reads, indicating that the ssDNA made by DRT1 *in vivo* is shorter than that made *in vitro*, regardless of bacteriophage infection. Consequently, cDNA complementary to the >70% adenosine-rich adducts (reads containing ≥70% thymidine) were not detected (**Fig. 5b; Extended Data Fig. 8b**). These data indicate that the A-rich ssDNA synthesized by DRT1 does not act as a template for DNA synthesis. While no discernable sequence motif was identified in the ≥70% adenosine reads, we observed that the base composition of purified DRT1 ssDNA determined via cDIP-seq matched closely to that determined for the same sample via LC nucleoside analysis, supporting the validity of our ≥70% adenosine filtering parameter (**Fig. 5c**). To test if ssDNA length increases upon bacteriophage T4 infection we conducted a western blot analysis of lysate from DRT1-expressing cells after bacteriophage infection. DRT1 migration was not altered relative to RT-inactive DRT1 (**Fig. 5d**), consistent with DRT1 ssDNA length remaining short *in vivo* and is not appreciably extended upon bacteriophage infection. The migration of DRT1 by SDS-PAGE correlates with the ssDNA length (**Extended Data Fig. 8c**). Together, these data support the function of DRT1 DNA adducts as structural elements rather than templates for toxic products.

**Fig. 5.**
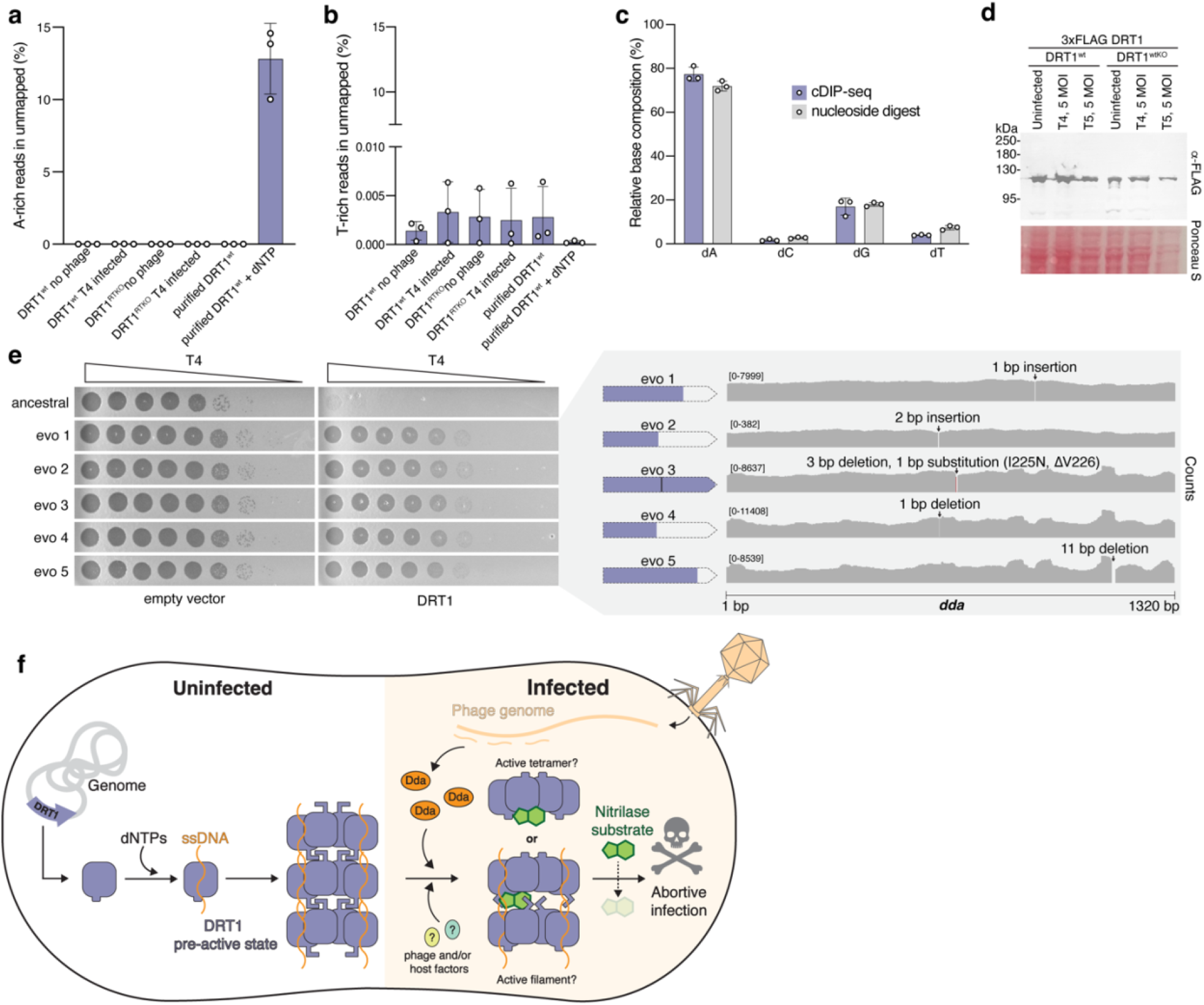
DNA adducts are a non-coding component of DRT1 defence. **a)** Percentage of unmapped reads containing greater than 70% adenine by cDIP-Seq. **b)** Percentage of unmapped reads containing greater than 70% thymidine by cDIP-Seq. **c)** Comparison of nucleotide composition by cDIP-Seq and LC nucleoside analysis of DNA synthesized by purified dNTP-reacted DRT1^wt^. **d)** Western blot of DRT1^wt^ and DRT1^RTKO^ pre, and post, infection with bacteriophage. **e)** Evolution of bacteriophage T4 against DRT1^wt^. Bacteriophage plaquing of five distinct evolutions after the 5th round. Genomic sequencing results of the evolved bacteriophage with zoomed in view of *dda* gene. **f)** Graphical model of DRT1 mechanism.

Phages often develop resistance to defence systems by acquiring mutations in elements that activate bacterial immune response. We serially passaged T4 and T5 phages on *E. coli* expressing DRT1 to isolate escape mutants. Bacteriophage T5 escapers were not observed despite repeated attempts. However, five T4 mutants with resistance to DRT1 were isolated (**Fig. 5e**). Whole-genome sequencing revealed that all five escapers acquired unique insertions or deletions within the T4 *dda* 5’→3’ single-stranded DNA helicase gene, all of which would result in mutated amino acids or truncated protein (**Fig. 5e**). To test if Dda protein is sufficient to trigger DRT1, the WT *dda* sequence was cloned into an arabinose-inducible vector and co-expressed with DRT1 or empty vector. Co-expression of DRT1 and Dda was not toxic *i*.*e*. Dda is necessary, but not sufficient, to activate DRT1 (**Extended Data Fig. 8d**). Indeed, of the fifteen bacteriophage sensitive to DRT1, twelve bacteriophage encode Dda or a Dda homologue (**Extended Data Fig. 8f**). All bacteriophage lacking Dda, or a homologue of Dda, are insensitive to DRT1. Given T4 single-stranded binding protein (SSB) is known to activate Dda^30^, we also tested the co-toxicity of SSB alone and dual expression with Dda. These data were inconclusive as SSB was toxic even in the absence of DRT1 (**Extended Data Fig. 8d**). To further probe for binding of Dda or other bacteriophage proteins to DRT1, we conducted pulldown assays using tagged DRT1 followed by unbiased proteomics analysis. Only two T4 proteins—Inh and Trna.2—were significantly enriched in DRT1^wt^ relative to the DRT1^RTKO^ control (**Extended Data Fig. 8e**). Given Inh and Trna.2 are only conserved in the four most closely related bacteriophage to T4 **(Extended Data Fig. 8f)** and these proteins appear not to have core metabolic function, we did not attempt further characterization of these hits. Therefore, the complete activation pathway of DRT1 remains elusive.

## Discussion

There is substantial risk when generating a contextually toxic immunity protein. Safety mechanisms are essential in preventing unintentional programmed cell death. Here we determined that DRT1 performs template-free, protein-primed, DNA synthesis to simultaneously activate and inhibit a nitrilase effector domain. Covalent DNA adducts stabilize tetrameric DRT1 which further assembles into filaments. The tetrameric nitrilase assembly renders activity^31,32^ while the filamentous structure induces C-terminal domain swapping to impose an inactive “closed-lid” conformation over the apo-nitrilase active site^28^. Opening and closing of a similarly conserved inter-protomeric active site lid is involved in the catalytic mechanism of other nitrilases^28,29^. At mid-stage of infection, a 5’→3’ single-stranded DNA helicase, along with other bacteriophage components, activate DRT1 (**Fig. 5f**). Our data demonstrate that non-templated DNA synthesis acts as both an activator and safety mechanism for DRT1 and may also serve as a sensor of bacteriophage replication enzymes.

Nitrilase enzymes are typified by supramolecular structure where homo-oligomeric assembly is a requirement for activity^31-34^. Consistent with this, RT-inactive DRT1−which is monomeric−is incapable of antiphage defence and is not toxic to the cell. It is reasonable to consider that DRT1, like DRT9, senses the nucleotide pool to enable selective formation of filaments^35,36^. Indeed, enzymes encoded by bacteriophage T4 and T5 responsible for dNTP biosynthesis upon infection are downregulated in the presence of DRT1. However, unlike DRT9 and the abortive infection polymerases, protein-primed DNA synthesis is not sufficient for DRT1 to impart immunity. DNA-adducts induce dormancy of DRT1 through nitrilase inhibition. We uncover no evidence to suggest that DRT1 DNA adducts perform any role beyond structural assembly, and perhaps sensing bacteriophage.

Most antiphage-defence systems with supramolecular structure assemble catalytically active filaments^37-41^. The pre-formed assembly of quiescent DRT1 prior to phage infection parallels the mechanism of retrons^12-14^, wherein the effectors are inactivated by a DNA cage within filaments in the absence of infection. In this sense, DRT1 exemplifies a minimal retron composed of a single polypeptide chain capable of generating an msDNA analogue to induce a pre-active state and potentially sense infection. It is enticing to speculate how Dda, and other phage components, may act upon DRT1-DNA adducts to induce an active state.

## Acknowledgments

We thank our colleagues C. Yancey, L. Ettwiller, P. Weigele, H. Lim, and D. Nye for helpful discussions. T. Blower and J. Eaglesham for comments on the manuscript. We thank S. Srikant (MIT) and M. Laub (MIT) for helpful discussions regarding bacteriophage evolution.

## Author contributions

Experiments were conceived and designed R.J.T., N.N., N.V.J., and J.S.M.. Bioinformatic analyses were performed by C.H.C. and S.R.J. Protein expression and purification was performed by N.N and R.J.T.. Nucleoside analysis was performed by N.D., I.R.C.J. and N.N.. Protein mass spectrometry data collection and analysis was performed by E.E.E. and A.H. Phage evolution, mass photometry, Western blotting, DRT reactivity, nuclease protection, cDIP-Seq, and RNA-Seq were performed by N.N. Phage spotting and liquid growth were performed by N.N. and A.N. Negative stain EM and cryo-EM data collection and analysis were performed by N.V.J. and J.S.M. Manuscript was prepared by N.N., N.V.J., and RJT, with support from all authors.

## Additional information

### Extended Data

#### Supplementary Information

The accession number for the DRT1 atomic model is 9YFD and density map EMD-72883.

#### Correspondence and requests for materials

Should be addressed to Robert J. Trachman or Jason S. McLellan

