## Extended Data for "Semirandom DNA adducts regulate a filamentous defence-associated reverse transcriptase"

### Materials and Methods

Unless otherwise stated, all reagents are from New England Biolabs (NEB). DRT genes were synthesized and cloned into pACYC184, pBAD or pET by GenScript. The sequences of the encoded proteins are listed in (**Extended Data Table S1**). All oligonucleotides were synthesized by Integrated DNA Technologies.

#### DRT phylogenetic reconstruction

The reverse transcriptase (RT) dataset was obtained from the supplementary material (xxxFileS1.nex) of Sharifi and Ye et al. [1]. The alignment was processed using Geneious Prime, and sequences were exported as an aligned FASTA file. Within the alignment, a close homolog of $DRT1^{wt}$, WP_075861988.1, which differs by only a single amino acid, was identified. This native sequence was replaced with $DRT1^{wt}$ to maintain the original alignment. This modified dataset was used to construct the phylogenetic tree using FastTree[2] with the LG[3] evolutionary model and discrete gamma model with ten rate categories. To improve accuracy, 4 rounds of subtree-prune-regraft (SPR) moves and slow nearest-neighbor interchanges (NNI) were applied [2] The tree was rooted at the midpoint and visualized using iTol (Interactive Tree of Life).

#### Phage defence plaque assays

All DRT systems were reconstituted in the surrogate host *E. coli* MG1655 from the low-copy plasmid pACYC184, wherein the DRT candidate gene has replaced the tetracycline resistance gene normally encoded in pACYC184 while maintaining the original pTet promoter. *E. coli* MG1655 was made competent using the Mix & Go kit (Zymo Research, T3001) and successful transformants were selected on LB agar containing 30 μg $mL^{-1}$ chloramphenicol (LB-cam). Several colonies were grown in 9 mL LB-cam overnight at 37 °C, and 200 μL of overnight culture was added to 6 mL of molten (51°C) top agar (1% soy peptone, 0.5% NaCl, 0.05% $MgCl_2 \cdot 6H_2O$, 10 mM $MgSO_4$, 0.75% agar, and 30 μg $mL^{-1}$ chloramphenicol), mixed, and poured on 10 cm LB-cam agar plates. Plates were incubated at 37° C for 5 hours, dried for 5 min under laminar flow, then inoculated with phages. Tenfold serial dilutions of phage were prepared immediately prior to assay setup in phage dilution (PD) buffer (50 mM Tris pH 7.5, 75 mM NaCl, and 10 mM $MgCl_2$). 3 μL of phage dilution was spotted, and the plates were incubated at 37° C overnight.

#### Liquid phage infection growth curve

Three different transformants of *E. coli* MG1655 harboring the indicated DRT plasmid were grown overnight in LB-cam. The cultures were diluted 1/100 in fresh LB-cam supplemented with 10 mM $MgSO_4$ and grown until OD600 ≈ 0.3, then diluted to a final $OD_{600}$ of 0.05 in the wells of a 96-well plate. Fresh dilutions of phage were prepared in LB-cam supplemented with 10 mM $MgSO_4$ and added to each well to yield a final MOI of 0.1, 1, or 10. Final well volume was 150 μL. Growth kinetics were monitored at 37 °C with shaking in a BioTek Synergy H1 (Agilent) plate reader with $OD_{600}$ reads every 5 min.

**Phage burst size assay**

To determine phage burst size and efficiency of center of infection, one-step growth curves were conducted as described with slight adaptation [4,5]. Overnight cultures of *E. coli* MG1655 harboring DRT1 in pACYC184 or empty vector were diluted 1/100 in LB-cam supplemented with 10 mM $MgSO_4$ then grown to OD600 ≈ 0.6. 2 mL of each culture was pelleted at 10,000g for 4 min, and resuspended in 900 µL of LB-cam supplemented with 10 mM $MgSO_4$. T4 phage was added to MOI = 0.0002 and allowed to adsorb for 10 min (no shaking) at 37°C. Unbound phage was removed by centrifuging at 10,000 x g for 2 min and resuspending in 300 µL fresh media, twice. 10 µL of the suspension was then added to 100 mL of pre-warmed LB-cam supplemented with 10 mM $MgSO_4$ and grown at 37°C with 200 rpm shaking. 1 mL aliquots were immediately withdrawn (15 min timepoint) and either directly plated for plaque enumeration (represents total phage in sample) or vortexed with 50 µL chloroform (represents free phage in sample) then plated. Every 10 min thereafter 1 mL aliquots were withdrawn and vortexed with 50 µL chloroform prior to plaque enumeration. PFU $mL^{-1}$ per initial infections = (average titer of free phages at late timepoints) / initial infections, where initial infections = 15 min timepoint titer before chloroform treatment) - (15 min timepoint titer after chloroform treatment).

**Cell survival following phage infection**

Overnight cultures of *E. coli* MG1655 harboring DRT1 in pACYC184 or empty vector were diluted 1/100 in LB-cam supplemented with 10 mM $MgSO_4$ then grown to OD600 = 0.4. Aliquots of 2 mL of each culture were mixed with the indicated MOI of T4 phage and incubated with shaking at 37°C for 15 min. Samples of 1 mL were then withdrawn and centrifuged at 10,000*g* for 3 min. The supernatant was removed, and the pellet was resuspended in 200 µL of LB. Resuspended cells were serially diluted in LB and plated on LB-cam plates. Plates were incubated at 37°C overnight prior to enumerating colonies.

**DRT1 expression and purification**

All recombinant DRT proteins were expressed from a custom pET vector called pMC009, which yields an N-terminal His14-MBP-SUMO fusion. Plasmids were transformed into T7 Express (NEB, C2566H) or One Shot BL21 Star (DE3) (Invitrogen, C601003). A colony from freshly transformed plates was used to inoculate 9 mL of lysogeny broth (LB) (1% soy peptone, 0.5% yeast extract, and 0.5% NaCl) supplemented with 50 µg $mL^{-1}$ kanamycin (LB-kan) and grown overnight at 37°C. This starter culture was diluted 100-fold into flasks containing 1 L of LB-kan, which were grown at 37° C until $OD_{600}$ reached approximately 0.6. Isopropyl-beta-D-thiogalactoside (IPTG) was then added to 0.35 mM and the cultures were incubated at 16°C overnight. Cultures were harvested by centrifugation at 4000 xg at 4°C. Cell pellets were resuspended in lysis buffer (50 mM Tris pH 8.0, 500 mM NaCl, 5 mM $MgCl_2$, 5 mM 2-mercaptoethanol, 2% glycerol and 0.1% Tween-20) supplemented with 1 x Halt protease inhibitor cocktail (Thermo, 78438). Cells were lysed by sonication for 3 min (10 sec on, 10 sec off) at 60% amplitude using a Q500 instrument (QSonica, Q500-110) on ice. Lysate was clarified by centrifugation, 0.22 µ filtered, then applied to three daisy-chained 5 mL MBPTrap HP columns (Cytiva, 28918779) equilibrated with lysis buffer. Protein was eluted in lysis buffer supplemented with 10 mM maltose. Fractions containing DRT were pooled and brought to 1 M NaCl final, followed by addition of 2 µM final concentration SENP1 SUMO protease and incubated overnight at 4°C for tag cleavage. The sample was then loaded in a 3.5 kDa cutoff bag (Spectrum Labs, 132726) and dialyzed against buffer containing 25 mM Tris pH 8.0, 1 M NaCl, 1 mM $MgCl_2$, 5 mM 2-mercaptoethanol, 0.1% Tween-20, and 50% glycerol. Dialysis was used to concentrate since we observed that DRT1aggregates upon Centricon filtration. Concentrated sample was purified via size exclusion chromatography (SEC) using a Superose 6 Increase 10/300 GL column (Cytiva, 29091596) equilibrated with SEC buffer (25 mM Tris pH 8.0, 250 mM NaCl, 1 mM $MgCl_2$, 5 mM 2-mercaptoethanol, and 2% glycerol). Fractions containing purified DRT protein were pooled, adjusted to 1 M NaCl final concentration, then mixed with 1 μM final SENP1 for a second round of tag cleavage overnight at 4°C. The sample was concentrated by dialysis against buffer containing 25 mM Tris pH 8.0, 1 M NaCl, 1 mM $MgCl_2$, 5 mM 2-mercaptoethanol, and 50% glycerol using the same 3.5 kDa cutoff membrane as above. Concentrated sample was loaded onto the Superose 6 Increase 10/300 GL column equilibrated with SEC buffer without glycerol (25 mM Tris pH 8.0, 250 mM NaCl, 1 mM $MgCl_2$, and 5 mM 2-mercaptoethanol). Pure fractions were pooled and dialyzed against 25 mM Tris pH 8.0, 250 mM NaCl, 1 mM $MgCl_2$, 5 mM 2-mercaptoethanol, and 50% glycerol in 3.5 kDa cutoff Slide-A-Lyzer cassettes (Thermo, 66333) to concentrate.

**DRT5S/L expression and purification**

DRT5S/L was expressed as described above for DRT1. Cell pellets were resuspended in lysis buffer (50 mM Tris pH 8.0, 500 mM NaCl, 5 mM $MgCl_2$, 5 mM 2-mercaptoethanol, 2% glycerol and 0.1% Tween-20) supplemented with 1 x Halt protease inhibitor cocktail (Thermo, 78438). Cells were lysed by sonication for 3 min (10s on, 10s off) at 60% amplitude using a Q500 instrument (QSonica, Q500-110) on ice. Lysate was clarified by centrifugation, 0.22 μ filtered, then applied to three daisy-chained 5 mL MBPTrap HP columns (Cytiva, 28918779) equilibrated with lysis buffer. Protein was eluted in lysis buffer supplemented with 10 mM maltose. The N-His-MBP tag was not cleaved.

**Intact DRT1 LC-MS analysis**

LC-MS was conducted using a Vanquish Flex UHPLC system coupled to an Orbitrap Eclipse Mass Spectrometer (Thermo Scientific, San Jose, CA, USA). Wild-type and dRT (100 fmol/μL) were reduced with 20 mM TCEP (Thermo Scientific, Waltham, MA) for 10 minutes at 37˚C in starting mobile phase conditions. Two picomoles (20 μL) was injected for separation on a 2.1x50mm PLRP-S analytical column (5μm, 1000Å, Agilent Technologies, Lexington, MA), using mobile phases: A) 0.1% difluoroacetic acid (DFA) in water, B) 0.1% DFA in acetonitrile, and a 10-minute gradient from 20-68% B at 0.2 mL/min. Mass spectra were acquired in low-pressure "Intact Protein" mode using a HESI source, ion funnel RF 80%, source fragmentation 20V, source voltage 3.8 kV, source and vaporizer temperatures of 300 °C, sheath gas 35 (a.u.) and aux gas 10 (a.u.). Mass spectra were collected at 7.5k resolution, AGC 1E5, with 50ms max injection time. UniDec deconvolution software [6,7] was used for intact mass determinations.

**Peptide LC-MS/MS using product ion triggered EThcD**

20 µg of DRT1 was digested according to the S-Trap micro spin column protocol (Protifi). Proteolytic digestion using 2 ug trypsin (New England Biolabs) was carried out at 37°C for 18 hours. Tryptic peptides were recovered by sequential elution with 0.2% formic acid in water, 50% acetonitrile, and 70% acetonitrile. The combined eluate was evaporated and reconstituted in 0.1% formic acid and 2% acetonitrile in water. After A205 quantification using a Nanodrop One Spectrophotometer (Thermo Scientific) the peptide concentration was adjusted to 200 ng/µL for LC-MS/MS analysis.

Peptides were separated on an EASY-nLC 1200 using an EASY-Spray C-18 analytical column (150 mm, 75 µm, 3 µm, 100 Å) coupled to an Orbitrap Eclipse mass spectrometer (Thermo Fisher Scientific). 200 ng of peptides were separated using the mobile phases: A) 0.1% formic acid in water, and B) 0.1% formic acid in 80% acetonitrile, and a 75 min linear gradient from 2% to 45% B at 300 nL/min. MS1 spectra were collected at 60k resolution at a 1.5-second cycle-time. MS/MS spectra were collected by data-dependent acquisition using HCD (25 NCE) with ion trap detection (1E5 AGC; *m/z* 110-2000), and a 10 second dynamic exclusion after two events (±10 ppm tolerance). An extended mass range of m/z 110-4000 was used for product-ion triggered EThcD, 30k orbitrap resolution using calibrated charge- dependent ETD reaction times, 5E5 AGC, and 25% supplemental HCD.

**Characterization of nucleotide-modified peptides**

Byonic (Protein Metrics, Cupertino, CA) was used to identify peptides and site-localize nucleotide modified amino acids using diagnostic fragment ions[8] (**Extended Data Table 3**). Spectra from an initial untargeted LC-MS/MS run were evaluated using Byonic MS/MS filtering to determine whether any contained the diagnostic fragments listed in (**Extended Data Table 3)**. If at least 2 diagnostic fragments for a given nucleotide were observed in the top 10 most abundant ions, the product-ion triggered EThcD LC-MS/MS method was implemented in a second analysis. After filtering by peptide spectral match quality (Byonic Score and PEP 2D score), the paired product-ion triggered HCD/EThcD MS/MS spectra were manually evaluated to confirm modifications at a specific amino acid residue.

**Mass photometry**

Samples were analyzed with a Refeyn TwoMP mass photometer on cation-coated slides (Refeyn, MP-CON-71002). Uncoated slides (Refeyn, MP-CON-41001) yielded similar results. The mass standard curve was constructed using the MassFerence P1 calibrant (Refeyn,MP-CON-41033) diluted into SEC buffer. Reaction mixtures contained 50 mM Tris pH 7.5, 150 mM NaCl, 5 mM $MgCl_2$, 2 mM dithiothreitol (DTT), approximately 0.5 mg $mL^{-1}$ DRT1 protein, and 0.1 mM dNTP mixture or individual ddNTP where indicated. This mixture was incubated for 15 min at 37°C, then diluted into SEC buffer to yield approximately 500 nM final coverslip concentration of DRT1.

**dNTP polymerization assays**

Template-independent DNA polymerization was assayed in reactions containing 50 mM Tris pH 7.5, 150 mM NaCl, 5 mM $MgCl_2$, 2 mM dithiothreitol (DTT), 0.4 mg $mL^{-1}$ RNAse A (NEB, T3018L) and approximately 0.5 mg $mL^{-1}$ DRT protein. This mixture was incubated for 15 min at 37°C to digest any contaminating RNA that might be present. DNA synthesis was then initiated by adding 0.1 mM of each dATP, dCTP, dGTP, and dTTP, unless otherwise indicated. Reactions were incubated at 37°C for 30 min. For SDS-PAGE analysis, reactions were quenched by mixing with SDS sample dye (NEB, B7703S) and heating at 95°C for 5 min, followed by electrophoresis on 4-20% Novex gel (Invitrogen, XP04200BOX) and staining with Coomassie blue.

For nucleic acid analysis, reactions were quenched by adding 25 mM EDTA and 0.4 mg $mL^{-1}$ proteinase K (NEB, P8107S) and incubating for an additional 15 min at 37°C to release DNA covalently attached to protein. To visualize products via electrophoresis, the reactions were mixed 1:1 with RNA Loading Dye (NEB, B0363S), heated at 65°C for 5 min, then resolved on 15% TBE-urea gels (Invitrogen, EC6885BOX). Gels were stained for 15 min with 1x SYBR Gold (Invitrogen, S11494) dissolved in 1x TBE buffer. Gels were rinsed briefly with distilled water then imaged under UV light. Low range ssRNA ladder (NEB, N0364S), in some cases mixed with microRNA marker (NEB, N2102S), was used to approximate the size of ssDNA products given the absence of a commercial ssDNA ladder in this size range.

**Nucleoside analysis liquid chromatography**

DRT reactions of 100 µL were prepared as described in the dNTP polymerization assays section. Nucleic acid was released by digestion with 0.4 mg $mL^{-1}$ proteinase K for 15 min at 37°C. The resulting sample was mixed with 700 µL of binding buffer and purified using the NEB Monarch spin PCR & DNA cleanup kit (T1130S). Purified DNA was eluted in water and then digested into nucleosides via the NEB nucleoside digestion mix (M0649S). For quantitative comparison of relative nucleoside composition, the raw integral corresponding to each nucleoside peak was divided by its respective extinction coefficient at 260 nm: A 15,200; C 7,050; G 12,010; and T 8,400 [9].

**Negative Stain Electron Microscopy**

Unreacted DRT1 samples or DRT1 reacted with dNTPs were diluted to 0.3 mg/mL in buffer containing 50 mM Tris pH 7.5, 150 mM NaCl, 5 mM MgCl2. Diluted samples were immediately applied to glow-discharged copper-supported carbon 400 mesh grids (Electron Microscopy Sciences) and stained with 2% methylamine tungstate (Nano-W, Nanoprobes). Grids were loaded onto either a JEOL NEOARM (200 kV) or JEOL1400F (120 kV) transmission electron microscope, each equipped with a OneView camera. Micrographs were acquired in 4 k x 4 k mode using Digital Micrograph (Gatan) with a nominal magnification of 50,000x (pixel size = 2.7 Å) on the NEOARM and 60,000x (pixel size = 3 Å) on the 1400F. Micrographs were exported to cryoSPARC for CTF correction, particle picking, and 2D classification. 3D volumes were generated using ab initio reconstruction, and data were further processed through heterogeneous and homogeneous refinements. Structural figures were produced using ChimeraX v1.8 [10].

**Cryo-EM sample preparation and data collection**

Sample containing 0.6 mg/mL DRT1 + dNTPs was applied to C-flat 1.2/1.3 300 mesh grids (Electron Microscopy Sciences) that had been glow discharged using a Leica EM ACE600 for 30 seconds with a 20 mA current. Using a Leica EM GP2 automatic plunging system set to 12 °C and 100% humidity, 4 uL of sample was applied to the backside of the grid and excess liquid was blotted away by contacting filter paper for 3 seconds. Sample application and blotting were repeated two times (3 blots total) before plunge-freezing the grid in liquid ethane.

10,188 movies were collected from a single grid using a Titan Krios TEM (Thermo Fisher) equipped with a K3 direct electron detector, Volta phase plate, and a Gatan imaging filter with a slit width of 20 eV. All movies were collected using SerialEM automation software[11]. Particles were imaged at a calibrated magnification of 0.413 Å/pixel, with an exposure of ~18 eps for 2.8s for a total exposure of ~50 e/$Å^2$. Additional details about data collection parameters can be found in **(Extended Data Table 2).**

**Cryo-EM data processing and structure building**

Motion correction and CTF estimation were performed using cryoSPARC v4.6.0 live processing. After all data was collected, pre-processed exposures were exported, and further processing was performed using cryoSPARC. Initial 2D classification was performed using 31 particles that were manually picked along the filamentous particle and extracted. The two resulting classes were used to guide additional particle picking using the filament tracer job with the filament diameter set to 140 Å, a separation distance of 0.85 diameters, and a minimum filament length of two diameters considered. The picks were then extracted and distributed into two classes containing particles. 176,480 particles were used to generate an ab initio reconstruction of three classes. The full particle stack containing 921,286 particles was carried into a heterogeneous refinement of the three classes, and the particles from the highest quality class were used for homogenous refinement of the best volume with no applied symmetry. Next, a helical refinement job was performed with no applied symmetry or helical parameters defined. The resulting volume was then analyzed using a symmetry search job set to search over a helical rise of 40-140 Å and helical twist of -180° – 180°. The search identified a helical order of 2, rise of 59.13 Å and twist of -121.22°. The 3D volume was then subjected to another round of helical refinement using these parameters and D2 point group symmetry. Local CTF refinement was performed on the refined volume and particle stack, and a final helical refinement was performed using non-uniform refinement with minimization over per-particle scale factors (input values, not reset to 1.0), including the defined helical parameters and applied D2 symmetry. An initial model of the complex was generated using AlphaFold 2 (https://alphafoldserver.com). The highest confidence output model was docked into the refined volume via ChimeraX v1.8 [10]. The structure was iteratively refined and completed using a combination of Phenix v1.21.2, Coot v0.9.2, and ISOLDE v1.8[12-14].

**Exonuclease protection assay**

DRT1 was first reacted with dNTPs to yield extended ssDNA adducts. Twenty microliter reactions containing 0.5 mg $mL^{-1}$ DRT1, 50 mM Tris pH 7.5, 150 mM NaCl, 5 mM $MgCl_2$, 2 mM DTT, 0.4 mg $mL^{-1}$ RNase A (NEB, T3018L), and 0.1 mM final concentration dNTP mix (NEB, N0447L) or an equivalent volume of water, were reacted for 15 min at 37°C. Next, 2.2 µL of ExoI reaction buffer and 1 µL ExoI (NEB, M0293L) or water were added to each reaction and incubated for 30 min at 37°C. Reactions were heated to 93°C for 10 min to inactivate ExoI. Reactions were then mixed with 1 µL of proteinase K (NEB, P8107S) and incubated for 30 min at 37°C. Samples were mixed with an equal volume of RNA loading dye, heated at 65°C for 5 min, then resolved on 15% TBE-urea gel followed by SYBR Gold staining.

**Illumina sequencing of DRT1 pulldown ssDNA**

The protocol for sequencing of ssDNA produced by DRT1 during phage infection was adapted from the cDIP-seq technique of Tang *et al* [15]. *E. coli* MG1655 was transformed with empty plasmid, or plasmid encoding N-MBP-tagged DRT1 WT or its D324N, D325N mutant. Colonies were inoculated into LB-cam, grown overnight at 37°C, then subcultured into 200 mL fresh LB-cam supplemented with 10 mM $MgSO_4$ and grown to $OD_{600}$ of 0.8. T4 phage, or an equivalent volume of phage buffer, was then added to yield MOI of 5 or 0. Cells were incubated with phage for 10 min at 37°C with 200 rpm shaking, then pelleted by centrifugation for 10 min at 10,000 x g at 4°C. The pellet was flash frozen on dry ice then stored at -80 °C.

Pellets were resuspended in 30 mL of pulldown lysis buffer (50 mM Tris pH 8.0, 250 mM NaCl, 5 mM MgCl2, 2 % glycerol, 0.1 % Tween 20, 2 mM DTT, 1x Halt inhibitor) and sonicated at 25% amplitude for 1 min total (2 s on, 10 s off). Cell debris was pelleted by centrifugation at 20,000g for 20 min at 4°C. The supernatants were mixed with 6 mL of 50% slurry amylose resin (NEB E8022L) that had been pre-washed 3x in pulldown lysis buffer. The lysates were rocked in 50 mL conical tubes with resin for 1 h at 4°C. The resin was gently pelleted by centrifugation for 1 min at 500g, supernatant discarded, then pellet resuspended in 30 mL of fresh pulldown lysis buffer. This was repeated twice more for a total of 3 washes. Protein was eluted by adding 1 mL of pulldown lysis buffer without Tween20 and supplemented with 50 mM maltose and rocking for 30 min at 4°C. Total protein in the eluates was quantified with detergent compatible Bradford assay (Thermo 23246).–Samples were normalized based on total protein, and 50 µL aliquots each containing approximately 20 µg of total protein were used for downstream purification.

Aliquots of eluate were mixed with 3 µL RNase A (NEB T3018L) and incubated at 37 °C for 15 min. Proteinase K (NEB, P8107S) was then added at 3 µL, and samples were digested for 1 h at 37 °C. The samples were purified using the ssDNA/RNA clean & concentrate kit (Zymo, D7011). Purified ssDNA was quantified via Qubit (Thermo Q10212). The control samples consisting of 20 µg of fully purified untagged DRT1 with or without *in vitro* reaction with 0.5 mM dNTPs were also subjected to the above steps.

Approximately 5 ng of ssDNA from each sample was used as starting material for the xGen ssDNA & Low-Input DNA Library Prep Kit (IDT 10009859). Library preparation followed the manufacturer's instructions with the following changes to SPRISelect bead volumes to maximize capture of short (< 200 nt) ssDNA. For post-extension cleanup, 1.8X SPRISelect bead volume was used and only a single bead purification step was done. For post-ligation cleanup, 1.4X SPRISelect bead volume was used. Indexing PCR used 12 cycles with xGen UDI primers (IDT 10005975). Post-PCR cleanup used 1X SPRISelect bead volume. Samples were sequenced on Illumina Miseq with 75 cycle single-end reads.

Trim Galore was used to remove adapter sequences and remove reads shorter than 10 bp. Reads were mapped to a reference file containing the MG1655 genome (NC_000913.3), T4 genome (GenBank: AF158101.6), and relevant plasmid sequences, using bwa with default settings. SAMtools (v 1.21) flagstat was used to analyze alignment statistics and count unmapped reads. SAMtools fasta was used to extract unmapped reads. A custom Python script was used to filter unmapped reads based on polyA or polyT content, with reads containing ≥ 70% A or ≥ 70% T being extracted for downstream analysis.

**Evolution of escape mutant phage**

Escape mutant phages were evolved as described [16]. For both phages T4 and T5, five independent phage populations were evolved against resistant host *E. coli* MG1655 pACYC-DRT1 in 96-well deep-well plates. In each plate, a control population was evolved with only the sensitive host containing empty pACYC plasmid. Overnight cultures in LB-cam were diluted 1/100, grown to an $OD_{600}$ of approximately 1.0, then diluted again to $OD_{600}$ of 0.00125 in either Teknova LB (L8000) supplemented with 1 mM $MgSO_4$ and 30 µg $mL^{-1}$ chloramphenicol for T4, or Teknova Minimal M9 Broth (M8000) supplemented with 30 µg $mL^{-1}$ chloramphenicol for T5. Each well of the plate was seeded with 200 µL of diluted bacteria. Each well was infected with 20 µL of tenfold serial dilutions of T4 or T5 in PD buffer, or 20 µL of PD buffer only to monitor contamination. The plates were sealed with breathable film and incubated at 37°C with 300 rpm shaking for 18 h. The plate was harvested by pooling the most diluted well that was fully cleared along with the first uncleared well for each population. The pooled samples were centrifuged at 4000 g for 20 min at 4°C to pellet cell debris. Supernatant was passed through 0.22 µm filters (Millipore UFC30GV00) and stored in tubes with 40 µL of chloroform added to prevent bacterial contamination.

**Evolved phage DNA extraction and Illumina sequencing**

To prepare for DNA extraction of evolved phages, each DRT1 evolution population and corresponding empty vector control population were plated on soft agar overlay of DRT1-expression or empty vector bacteria, respectively. A single plaque from each sample was used to infect 10 mL cultures of the same bacterial host grown to $OD_{600}$ of 0.5 in LB-cam supplemented with 10 mM $MgSO_4$. Cultures were grown for 5 h at 37°C to allow phage propagation. The cultures were centrifuged at 4,000g for 20 min at 4°C, and supernatant was decanted into fresh tubes containing 1 g PEG 8000 (10% final) and 0.58 g NaCl (1M final). Tubes were mixed by inversion until dissolved then incubated at 4 °C overnight. Tubes were centrifuged at 4000 g for 30 min to pellet phages, which were then resuspended in 500 µL of 5 mM $MgSO_4$. Each sample received 1.25 µL of DNase1 (NEB, M0303S) and 1.25 µL of RNAse A (NEB, T3018L) followed by incubation at 37°C for 1 h. Each sample then received 30 µL of 0.5 M EDTA pH 8.0 (30 mM final), 25 µL of 10% SDS (0.5% final), and 1.25 µL of proteinase K (NEB, P8107S) (20 µg total) followed by 1h incubation at 60 °C. Samples were cooled to room temperature then mixed with an equal volume of phenol:chloroform (1:1). Following centrifugation for 10 min at 10,000 g, the supernatant was transferred to fresh tubes and phenol:chloroform extraction was repeated. An equal volume of chloroform was mixed with the supernatant and centrifuged for 10 min at 10,000 g. Supernatant was transferred to fresh tubes and mixed with a 1/10 volume of 3M sodium acetate pH 5.2 and 2.5 volumes of ice cold 100% ethanol. The mixture was incubated on ice for 30 min to precipitate DNA, which was then pelleted by centrifugation for 20 min at 16, 000g and 4°C. The DNA pellet was washed twice with 500 µL 70% ethanol then dried with tube caps open for 15 min. DNA was resuspended in 0.1x TE buffer (1 mM Tris-HCl pH 8.0, 0.1 mM EDTA).

Purified phage DNA was diluted to 50 ng $\mu L^{-1}$ and sheared using a Covaris ML230 in 8-AFA tubes (Covaris, 520292) to yield DNA fragments of approximately 175 bp. Sheared DNA was then prepared for sequencing using the NEBNext Ultra II library prep kit for Illumina (NEB, E7645S) and NEBNext multiplex oligos (NEB, E6446S). Agilent Tapestation was used to verify library quality and concentrations. Libraries were sequenced via Illumina NextSeq500 using 150 bp paired end reads.

Reads were aligned to the reference T4 genome (GenBank: AF158101.6) using BWA and Samtools. The consensus assembly of each sample was then extracted using Samtools. The extracted assembly of the DRT1 evolution clones was then aligned to the extracted assembly of the empty plasmid control evolution in Geneious Prime 2024-10-19. Coverage tracks were visualized in IGV.

**Bacteriophage Phylogenetic tree**

Viral phylogenetic analysis was carried out by the VICTOR web service (https://ggdc.dsmz.de/victor.php) [17]. Pairwise comparisons of the amino acid sequences were conducted using the Genome-BLAST Distance Phylogeny (GBDP) method [18] under settings recommended for prokaryotic viruses [17]. The resulting intergenomic distances were used to infer a balanced minimum evolution tree with branch support via FASTME including SPR postprocessing [19] for using the D6 distance formula. Branch support was inferred from 100 pseudo-bootstrap replicates each. Trees were rooted at the midpoint and visualized with ggtree [20]. Taxon boundaries at the species, genus and family level were estimated with the OPTSIL program [21], the recommended clustering thresholds [17] and an F value (fraction of links required for cluster fusion) of 0.5 [22]. The tree was rooted at the midpoint and visualized using iTOL (https://academic.oup.com/nar/article/52/W1/W78/7645242).

**Homolog identification**

Structures were predicted for all proteins annotated in the 23 phage genomes, using AlphaFold2 [23] through ColabFold [24]. Predicted structures for T4 Dda (UniProt: P32270), Inh (UniProt: P18058), and Trna.2 (UniProt: P13324) were used as queries in a Reseek [25] search, in 'sensitive' mode, against a database of the phage predicted structures. Hits with a Reseek Alignment Quality (AQ) score above 0.75 were considered as homologs.

**Whole transcriptome sequencing**

*E. coli* MG1655 transformed with DRT constructs in pACYC184 was grown overnight in LB-cam at 37°C. The bacteria were diluted 100-fold into fresh LB-cam supplemented with 10 mM final $MgSO_4$ and grown at 37°C to $OD_{600}$ of approximately 0.25. T4 phage was then added at 1 MOI, which was calculated assuming $8x10^8$ bacterial cells in 1 mL culture at $OD_{600}$ of 1.0. T5 phage was added at 5 MOI. Samples of 1 mL volume were dispensed into the wells of a 24-well plate. The plate was incubated at 37° C for 30 min with 240 rpm shaking, then centrifuged for 10 min at 3,500 x g and 4°C to pellet cells. The supernatant was discarded and pellets frozen on dry ice. Thawed pellets were resuspended in 25 µL TBS (50 mM Tris pH 7.0, 150 mM NaCl) and 10 µL T4 lysozyme (NEB, P8115L) and incubated for 5 min at 25° C. The RNA was then purified according to the manufacturer's specifications for enzymatic lysis using the Monarch Total RNA Miniprep Kit (NEB, T2010S), beginning with addition of 300 µL of RNA Lysis Buffer to each sample after lysozyme. The RNA integrity number was verified to be above 7 for all samples via Agilent Tapestation. 500 ng of total RNA per sample was then depleted of ribosomal RNA using the NEBNext rRNA Depletion Kit (Bacteria) (NEB, E7850L). cDNA libraries were produced using the NEBNext Ultra II Directional RNA Library Prep Kit for Illumina (NEB, E7760S) with PCR amplification via single index NEBNext Multiplex Oligos (NEB, E7335S and E7730S). Library quality and concentrations were verified with Agilent Tapestation. Libraries were sequenced via Illumina NextSeq500 using 75 bp paired end reads.

Transcriptomic data were processed using the following tools in the Galaxy server and R. Adapters were trimmed with Trim Galore using default parameters, followed by quality control using

FastQC. Reads were mapped to either the T4 phage genome (GenBank: AF158101.6) or T5 phage genome (NCBI: NC_005859.1) using HISAT2 with strandedness set to reverse (RF). Mapped reads were counted using htseq-count with strandedness set to reverse. Differential gene expression was determined using DESeq2 with independent filtering and outlier filtering turned off.

**Western blotting**

Overnight cultures of *E. coli* MG1655 transformed with pACYC184 encoding C-terminal 3xFLAG-tagged DRT1 were diluted 100-fold into fresh LB-cam supplemented with 10 mM final $MgSO_4$ and grown at 37°C to $OD_{600}$ of approximately 0.3, at which point 5 MOI of phage (or equal volume of phage dilution buffer) was added. The cultures were then incubated at 37°C for 20 min and harvested by centrifugation at 4000g for 10 min. Pellets were resuspended in 500 µL of IP lysis buffer (20 mM Tris-HCl, pH 7.5, 150 mM KCl, 2 mM $MgCl_2$, 0.2% Triton X-100, and 1x Halt protease inhibitor), then lysed via sonication (20% amplitude for 1 min 30 s, 2 s on 5 s off). Debris was cleared by centrifugation at 10,000g for 5 min. Supernatant was mixed with SDS loading dye, heated at 95°C for 5 min, then run on 4-20% gel (Novex XP04205BOX) at 200 V for 1 h. Proteins were transferred to PVDF membrane (Biorad, 1620260) using the BioRad Turbo Transfer system. The membranes were washed 3x with distilled water, stained with Ponceau, briefly destained in distilled water, then dried on filter paper at room temperature for 1 h. Membranes were blocked by rocking with 5% milk in TBST at 4°C overnight. Primary mouse anti-FLAG antibody (Sigma, F1804) was added to the blocking solution at 1/1000 and rocked for 2 hours at 4°C. The membranes were washed 3x in TBST, then incubated with DyLight 800 goat anti-mouse 4xPEG conjugate secondary antibody (Invitrogen, SA5-35521) in TBST at 1/100,000 for 1hour at room temperature. After washing 3x in TBST, the membranes were imaged with a Licor Odyssey.

**Pulldown and mass spectrometry**

Protein samples for pulldown analysis were prepared as described in the '**Illumina sequencing of DRT1 pulldown ssDNA section**' above. Biological triplicates of each sample type were reduced, alkylated, digested, and chromatographically separated using the methods outlined in the '**Peptide LC-MS/MS using product ion triggered EThcD section**' above.

For this analysis, an Orbitrap Exploris Mass Spectrometer (Thermo Fisher Scientific) was used, equipped with a FAIMS Pro Interface (Thermo Fisher Scientific). A 160 min linear gradient was used from 2% to 35% B at 300 nL/min. The FAIMS Pro module was used to further separate peptides by cycling compensation voltage (CV) between -50 and -70V. For each CV, MS1 spectra spanning *m/z* 400-1450 were collected at 120k resolution. MS/MS spectra were collected by data-dependent acquisition using HCD (30 NCE, 45k resolution) at a 1.5-second cycle-time (2E5 AGC), a 45-second dynamic exclusion after a single event (±10 ppm tolerance), and an automatic *m/z* scan range with a 1.2 *m/z* isolation window. Raw data were searched using two reference proteomes and a purified protein FASTA: the E. coli K-12 strain MG1655 proteome, T4 phage proteome, and MBP-DRT1 sequences. Raw LC-MS/MS were analyzed using the LQF-MBR workflow within FragPipe (FragPipe v22.0; MSFragger v4.1; IonQuant v1.10.27; Philosopher v5.11; Python 3.11.11). Results were processed using Fragpipe Analyst v1.18 using an intensity-based DDA LFQ approach [26] and an all-pairs approach to compare each experimental condition with Perseus-type imputation and Benjamini Hochberg FDR correction [27].

### Coexpression toxicity assays

Toxicity of co-expressing DRT1 with candidate phage trigger proteins was assayed as described [28]. Briefly, single colonies of *E. coli* MG1655 harboring pACYC184-DRT1or empty pACYC184 plus pBAD encoding the indicated phage protein were grown for 6 h at 37°C in LB-glucose to saturation. A 200 µL aliquot of each culture was pelleted at 4,000*g* for 10 min, washed once in 1× PBS and resuspended in 400 µl of 1× PBS. The cells were then serially diluted tenfold in 1× PBS and 3 µL was spotted on plates composed of M9 medium (6.4 g $l^{-1}$ $Na_2HPO_4$-$7H_2O$, 1.5 g $l^{-1}$ $KH_2PO_4$, 0.25 g $l^{-1}$ NaCl, 0.5 g $l^{-1}$ $NH_4Cl$) supplemented with 0.1% casamino acids, 0.4% glycerol, 2 mM $MgSO_4$, 0.1 mM $CaCl_2$, 5% Teknova LB (v/v), and 1.5% agar (w/v). Plates were incubated overnight at 37 °C.

### Extended Data Table 1. Sequences used in this study

| Protein | Amino acid sequence |
|---|---|
| $DRT1^{WT}$ | MKLLNKKYYNLEPNFDYLKDSFILGLAWKKTDRFVRTHNWYADLLALDKCAFDISDEVTSWSSEAVNRNLSKSDIELIPAPKRASWFFNKGKWTTNKDDRKLRPLANISIKDQS<br>FATAVTMCLADAIETRQKDCSLRNLGYAEHVNKKVVSYGNRLVCDWDNERARFRWGGSEYYRKFSTDYRNFLQRPVYIGRETVNKISEIDDVYIISLDLKNFFGSIKISTLLEKLK<br>KISSDHYDHNIINENEFWGLASQILNWEWPEDSLSLLKNLDIEEENVGLPQGLASAGALANAYLIEFDESLTSNLRTKIDGSQIVLHDYCRYVDDIRLVISGEALTNEEIKKSIHEL<br>VQRILDETLDQDESDNEPYLKINDKKTYILGLSDIDNGSSLTNRINEIQNEVGTSSIPERNGLDNNIPALQQLLLTEQDNFLEDADGLFSGFNNDKSIKLESLRRFSAHRLETSLA<br>NKSKLISPAERKQFDNESALIAKKLLKTWLKDPSIMVIFRKAITINPNLDAYKTILEIIFLRIQHNREKCDRYIMLYLLSDIFRSVVDIYRKLSPECTSDYQKLMSEVTSFAHKLLSCRS<br>VIPNYAYQQALFYLAVINKPFIASKKSSSDLSKLQHVLIKRHLEPLSSSDGYLFELSAQISKDYQANVAFLLSHTNTTEVVDSIVEKFAYRGGEFWNSIWKEFVRTKDKVRINKFRW<br>AIPKNESKPNGSDHYLSSVISFKENPFKYEHALLKLGIALVNLLEDTEKNVWQSDGKQYSPHEIKVKLEGHSTSWTELWRQNSIISCSIDKIAQDPRYESPKWLVNCQQAKNDE<br>KKIYWVCSVLRSAALGNVDYTQRNDLKPDRVEYDGIHSQFYKRRMGMLHTPESIVGSYATITDWFASFLQHGLQWPGFSSSYINQEDILSITSLGEFKKCLLERLGYLNTQICTS<br>SSVPTLPTVVNRPELASNYFRIVTVQQLFPKDKHFHPSDVTLDNPEVRWKHREHLAEICKLTEQTLDAKLKTESRDHTSTADLIVFSELAVHPED EDIVRALAFRTRSIIFCGFVFC<br>EQDGQIVNKARWIIPDSSESGTQWRVRDQGKFHMTSGEMSLGVHGYRPSQHIISIEGHTEGPFKLTGAICYDATDIKLAADLRDLTDMFVIAAYNKDVDTFDNMASALQWHM<br>YQHVIITNTGEYGGSTMQAPYKEKHHKLISHAHGTGQIAISTADIDLAAFRRKIKEYKKTKTQPAGFKRKH |
| $DRT1^{RT\ KO}$ (D324N, D325N) | MKLLNKKYYNLEPNFDYLKDSFILGLAWKKTDRFVRTHNWYADLLALDKCAFDISDEVTSWSSEAVNRNLSKSDIELIPAPKRASWFFNKGKWTTNKDDRKLRPLANISIKDQS<br>FATAVTMCLADAIETRQKDCSLRNLGYAEHVNKKVVSYGNRLVCDWDNERARFRWGGSEYYRKFSTDYRNFLQRPVYIGRETVNKISEIDDVYIISLDLKNFFGSIKISTLLEKLK<br>KISSDHYDHNIINENEFWGLASQILNWEWPEDSLSLLKNLDIEEENVGLPQGLASAGALANAYLIEFDESLTSNLRTKIDGSQIVLHDYCRYVNNIRLVISGEALTNEEIKKSIHEL<br>VQRILDETLDQDESDNEPYLKINDKKTYILGLSDIDNGSSLTNRINEIQNEVGTSSIPERNGLDNNIPALQQLLLTEQDNFLEDADGLFSGFNNDKSIKLESLRRFSAHRLETSLA<br>NKSKLISPAERKQFDNESALIAKKLLKTWLKDPSIMVIFRKAITINPNLDAYKTILEIIFLRIQHNREKCDRYIMLYLLSDIFRSVVDIYRKLSPECTSDYQKLMSEVTSFAHKLLSCRS<br>VIPNYAYQQALFYLAVINKPFIASKKSSSDLSKLQHVLIKRHLEPLSSSDGYLFELSAQISKDYQANVAFLLSHTNTTEVVDSIVEKFAYRGGEFWNSIWKEFVRTKDKVRINKFRW<br>AIPKNESKPNGSDHYLSSVISFKENPFKYEHALLKLGIALVNLLEDTEKNVWQSDGKQYSPHEIKVKLEGHSTSWTELWRQNSIISCSIDKIAQDPRYESPKWLVNCQQAKNDE<br>KKIYWVCSVLRSAALGNVDYTQRNDLKPDRVEYDGIHSQFYKRRMGMLHTPESIVGSYATITDWFASFLQHGLQWPGFSSSYINQEDILSITSLGEFKKCLLERLGYLNTQICTS<br>SSVPTLPTVVNRPELASNYFRIVTVQQLFPKDKHFHPSDVTLDNPEVRWKHREHLAEICKLTEQTLDAKLKTESRDHTSTADLIVFSELAVHPED EDIVRALAFRTRSIIFCGFVFC<br>EQDGQIVNKARWIIPDSSESGTQWRVRDQGKFHMTSGEMSLGVHGYRPSQHIISIEGHTEGPFKLTGAICYDATDIKLAADLRDLTDMFVIAAYNKDVDTFDNMASALQWHM<br>YQHVIITNTGEYGGSTMQAPYKEKHHKLISHAHGTGQIAISTADIDLAAFRRKIKEYKKTKTQPAGFKRKH |
| $DRT1^{Nit\ KO}$ (C1116A) | MKLLNKKYYNLEPNFDYLKDSFILGLAWKKTDRFVRTHNWYADLLALDKCAFDISDEVTSWSSEAVNRNLSKSDIELIPAPKRASWFFNKGKWTTNKDDRKLRPLANISIKDQS<br>FATAVTMCLADAIETRQKDCSLRNLGYAEHVNKKVVSYGNRLVCDWDNERARFRWGGSEYYRKFSTDYRNFLQRPVYIGRETVNKISEIDDVYIISLDLKNFFGSIKISTLLEKLK<br>KISSDHYDHNIINENEFWGLASQILNWEWPEDSLSLLKNLDIEEENVGLPQGLASAGALANAYLIEFDESLTSNLRTKIDGSQIVLHDYCRYVDDIRLVISGEALTNEEIKKSIHEL<br>VQRILDETLDQDESDNEPYLKINDKKTYILGLSDIDNGSSLTNRINEIQNEVGTSSIPERNGLDNNIPALQQLLLTEQDNFLEDADGLFSGFNNDKSIKLESLRRFSAHRLETSLA<br>NKSKLISPAERKQFDNESALIAKKLLKTWLKDPSIMVIFRKAITINPNLDAYKTILEIIFLRIQHNREKCDRYIMLYLLSDIFRSVVDIYRKLSPECTSDYQKLMSEVTSFAHKLLSCRS<br>VIPNYAYQQALFYLAVINKPFIASKKSSSDLSKLQHVLIKRHLEPLSSSDGYLFELSAQISKDYQANVAFLLSHTNTTEVVDSIVEKFAYRGGEFWNSIWKEFVRTKDKVRINKFRW<br>AIPKNESKPNGSDHYLSSVISFKENPFKYEHALLKLGIALVNLLEDTEKNVWQSDGKQYSPHEIKVKLEGHSTSWTELWRQNSIISCSIDKIAQDPRYESPKWLVNCQQAKNDE<br>KKIYWVCSVLRSAALGNVDYTQRNDLKPDRVEYDGIHSQFYKRRMGMLHTPESIVGSYATITDWFASFLQHGLQWPGFSSSYINQEDILSITSLGEFKKCLLERLGYLNTQICTS<br>SSVPTLPTVVNRPELASNYFRIVTVQQLFPKDKHFHPSDVTLDNPEVRWKHREHLAEICKLTEQTLDAKLKTESRDHTSTADLIVFSELAVHPED EDIVRALAFRTRSIIFCGFVFC<br>EQDGQIVNKARWIIPDSSESGTQWRVRDQGKFHMTSGEMSLGVHGYRPSQHIISIEGHTEGPFKLTGAIAYDATDIKLAADLRDLTDMFVIAAYNKDVDTFDNMASALQWHM<br>YQHVIITNTGEYGGSTMQAPYKEKHHKLISHAHGTGQIAISTADIDLAAFRRKIKEYKKTKTQPAGFKRKH |
| DRT1 T401A | MKLLNKKYYNLEPNFDYLKDSFILGLAWKKTDRFVRTHNWYADLLALDKCAFDISDEVTSWSSEAVNRNLSKSDIELIPAPKRASWFFNKGKWTTNKDDRKLRPLANISIKDQS<br>FATAVTMCLADAIETRQKDCSLRNLGYAEHVNKKVVSYGNRLVCDWDNERARFRWGGSEYYRKFSTDYRNFLQRPVYIGRETVNKISEIDDVYIISLDLKNFFGSIKISTLLEKLK<br>KISSDHYDHNIINENEFWGLASQILNWEWPEDSLSLLKNLDIEEENVGLPQGLASAGALANAYLIEFDESLTSNLRTKIDGSQIVLHDYCRYVDDIRLVISGEALTNEEIKKSIHEL<br>VQRILDETLDQDESDNEPYLKINDKKTYILGLSDIDNGSSLTNRINEIQNEVGASSIPERNGLDNNIPALQQLLLTEQDNFLEDADGLFSGFNNDKSIKLESLRRFSAHRLETSLA<br>NKSKLISPAERKQFDNESALIAKKLLKTWLKDPSIMVIFRKAITINPNLDAYKTILEIIFLRIQHNREKCDRYIMLYLLSDIFRSVVDIYRKLSPECTSDYQKLMSEVTSFAHKLLSCRS<br>VIPNYAYQQALFYLAVINKPFIASKKSSSDLSKLQHVLIKRHLEPLSSSDGYLFELSAQISKDYQANVAFLLSHTNTTEVVDSIVEKFAYRGGEFWNSIWKEFVRTKDKVRINKFRW<br>AIPKNESKPNGSDHYLSSVISFKENPFKYEHALLKLGIALVNLLEDTEKNVWQSDGKQYSPHEIKVKLEGHSTSWTELWRQNSIISCSIDKIAQDPRYESPKWLVNCQQAKNDE |

| | |
|---|---|
| | KKIYWVCSVLRSAALGNVDYTQRNDLKPDRVEYDGIHSQFYKRRMGMLHTPESIVGSYATITDWFASFLQHGLQWPGFSSSYINQEDILSITSLGEFKKCLLERLGYLNTQICTS<br>SSVPTLPTVVNRPELASNYFRIVTVQQLFPKDKHFHPSDVTLDNPEVRWKHREHLAEICKLTEQTLDAKLKTESRDHTSTADLIVFSELAVHPEDEDIVRALAFRTRSIIFCGFVFC<br>EQDGQIVNKARWIIPDSSESGTQWRVRDQGKFHMTSGEMSLGVHGYRPSQHIISIEGHTEGPFKLTGAICYDATDIKLAADLRDLTDMFVIAAYNKDVDTFDNMASALQWHM<br>YQHVIITNTGEYGGSTMQAPYKEKHHKLISHAHGTGQIAISTADIDLAAFRRKIKEYKKTKTQPAGFKRKH |
| DRT1<br>S402A | MKLLNKKYYNLEPNFDYLKDSFILGLAWKKTDRFVRTHNWYADLLALDKCAFDISDEVTSWSSEAVNRNLSKSDIELIPAPKRASWFFNKGKWTTNKDDRKLRPLANISIKDQS<br>FATAVTMCLADAIETRQKDCSLRNLGYAEHVNKKVVSYGNRLVCDWDNERARFRWGGSEYYRKFSTDYRNFLQRPVYIGRETVNKISEIDDVYIISLDLKNFFGSIKISTLLEKLK<br>KISSDHYDHNIINENEFWGLASQILNWEWPEDSLSLLKNLDIEEENVGLPQGLASAGALANAYLIEFDESLTSNLRTKIDGSQIVLHDYCRYVDDIRLVISGEALTNEEIKKSIHEL<br>VQRILDETLDQDESDNEPYLKINDKKTYILGLSDIDNGSSLTNRINEIQNEVGTASIPERNGLDNNIPALQQLLLTEQDNFLEDADGLFSGFNNDKSIKLESLRRFSAHRLETSLA<br>NKSKLISPAERKQFDNESALIAKKLLKTWLKDPSIMVIFRKAITINPNLDAYKTILEIIFLRIQHNREKCDRYIMLYLLSDIFRSVVDIYRKLSPECTSDYQKLMSEVTSFAHKLLSCRS<br>VIPNYAYQQALFYLAVINKPFIASKKSSSDLSKLQHVLIKRHLEPLSSSDGYLFELSAQISKDYQANVAFLLSHTNTTEVVDSIVEKFAYRGGEFWNSIWKEFVRTKDKVRINKFRW<br>AIPKNESKPNGSDHYLSSVISFKENPFKYEHALLKLGIALVNLLEDTEKNVWQSDGKQYSPHEIKVKLEGHSTSWTELWRQNSIISCSIDKIAQDPRYESPKWLVNCQQAKNDE<br>KKIYWVCSVLRSAALGNVDYTQRNDLKPDRVEYDGIHSQFYKRRMGMLHTPESIVGSYATITDWFASFLQHGLQWPGFSSSYINQEDILSITSLGEFKKCLLERLGYLNTQICTS<br>SSVPTLPTVVNRPELASNYFRIVTVQQLFPKDKHFHPSDVTLDNPEVRWKHREHLAEICKLTEQTLDAKLKTESRDHTSTADLIVFSELAVHPEDEDIVRALAFRTRSIIFCGFVFC<br>EQDGQIVNKARWIIPDSSESGTQWRVRDQGKFHMTSGEMSLGVHGYRPSQHIISIEGHTEGPFKLTGAICYDATDIKLAADLRDLTDMFVIAAYNKDVDTFDNMASALQWHM<br>YQHVIITNTGEYGGSTMQAPYKEKHHKLISHAHGTGQIAISTADIDLAAFRRKIKEYKKTKTQPAGFKRKH |
| DRT1<br>S403A | MKLLNKKYYNLEPNFDYLKDSFILGLAWKKTDRFVRTHNWYADLLALDKCAFDISDEVTSWSSEAVNRNLSKSDIELIPAPKRASWFFNKGKWTTNKDDRKLRPLANISIKDQS<br>FATAVTMCLADAIETRQKDCSLRNLGYAEHVNKKVVSYGNRLVCDWDNERARFRWGGSEYYRKFSTDYRNFLQRPVYIGRETVNKISEIDDVYIISLDLKNFFGSIKISTLLEKLK<br>KISSDHYDHNIINENEFWGLASQILNWEWPEDSLSLLKNLDIEEENVGLPQGLASAGALANAYLIEFDESLTSNLRTKIDGSQIVLHDYCRYVDDIRLVISGEALTNEEIKKSIHEL<br>VQRILDETLDQDESDNEPYLKINDKKTYILGLSDIDNGSSLTNRINEIQNEVGTSAIPERNGLDNNIPALQQLLLTEQDNFLEDADGLFSGFNNDKSIKLESLRRFSAHRLETSLA<br>NKSKLISPAERKQFDNESALIAKKLLKTWLKDPSIMVIFRKAITINPNLDAYKTILEIIFLRIQHNREKCDRYIMLYLLSDIFRSVVDIYRKLSPECTSDYQKLMSEVTSFAHKLLSCRS<br>VIPNYAYQQALFYLAVINKPFIASKKSSSDLSKLQHVLIKRHLEPLSSSDGYLFELSAQISKDYQANVAFLLSHTNTTEVVDSIVEKFAYRGGEFWNSIWKEFVRTKDKVRINKFRW<br>AIPKNESKPNGSDHYLSSVISFKENPFKYEHALLKLGIALVNLLEDTEKNVWQSDGKQYSPHEIKVKLEGHSTSWTELWRQNSIISCSIDKIAQDPRYESPKWLVNCQQAKNDE<br>KKIYWVCSVLRSAALGNVDYTQRNDLKPDRVEYDGIHSQFYKRRMGMLHTPESIVGSYATITDWFASFLQHGLQWPGFSSSYINQEDILSITSLGEFKKCLLERLGYLNTQICTS<br>SSVPTLPTVVNRPELASNYFRIVTVQQLFPKDKHFHPSDVTLDNPEVRWKHREHLAEICKLTEQTLDAKLKTESRDHTSTADLIVFSELAVHPEDEDIVRALAFRTRSIIFCGFVFC<br>EQDGQIVNKARWIIPDSSESGTQWRVRDQGKFHMTSGEMSLGVHGYRPSQHIISIEGHTEGPFKLTGAICYDATDIKLAADLRDLTDMFVIAAYNKDVDTFDNMASALQWHM<br>YQHVIITNTGEYGGSTMQAPYKEKHHKLISHAHGTGQIAISTADIDLAAFRRKIKEYKKTKTQPAGFKRKH |
| DRT1<br>K90G,<br>K92E | MKLLNKKYYNLEPNFDYLKDSFILGLAWKKTDRFVRTHNWYADLLALDKCAFDISDEVTSWSSEAVNRNLSKSDIELIPAPKRASWFFNGGEWTTNKDDRKLRPLANISIKDQ<br>SFATAVTMCLADAIETRQKDCSLRNLGYAEHVNKKVVSYGNRLVCDWDNERARFRWGGSEYYRKFSTDYRNFLQRPVYIGRETVNKISEIDDVYIISLDLKNFFGSIKISTLLEKL<br>KKISSDHYDHNIINENEFWGLASQILNWEWPEDSLSLLKNLDIEEENVGLPQGLASAGALANAYLIEFDESLTSNLRTKIDGSQIVLHDYCRYVDDIRLVISGEALTNEEIKKSIHE<br>LVQRILDETLDQDESDNEPYLKINDKKTYILGLSDIDNGSSLTNRINEIQNEVGTSSIPERNGLDNNIPALQQLLLTEQDNFLEDADGLFSGFNNDKSIKLESLRRFSAHRLETSL<br>ANKSKLISPAERKQFDNESALIAKKLLKTWLKDPSIMVIFRKAITINPNLDAYKTILEIIFLRIQHNREKCDRYIMLYLLSDIFRSVVDIYRKLSPECTSDYQKLMSEVTSFAHKLLSCR<br>SVIPNYAYQQALFYLAVINKPFIASKKSSSDLSKLQHVLIKRHLEPLSSSDGYLFELSAQISKDYQANVAFLLSHTNTTEVVDSIVEKFAYRGGEFWNSIWKEFVRTKDKVRINKFR<br>WAIPKNESKPNGSDHYLSSVISFKENPFKYEHALLKLGIALVNLLEDTEKNVWQSDGKQYSPHEIKVKLEGHSTSWTELWRQNSIISCSIDKIAQDPRYESPKWLVNCQQAKN<br>DEKKIYWVCSVLRSAALGNVDYTQRNDLKPDRVEYDGIHSQFYKRRMGMLHTPESIVGSYATITDWFASFLQHGLQWPGFSSSYINQEDILSITSLGEFKKCLLERLGYLNTQI<br>CTSSSVPTLPTVVNRPELASNYFRIVTVQQLFPKDKHFHPSDVTLDNPEVRWKHREHLAEICKLTEQTLDAKLKTESRDHTSTADLIVFSELAVHPEDEDIVRALAFRTRSIIFCGF<br>VFCEQDGQIVNKARWIIPDSSESGTQWRVRDQGKFHMTSGEMSLGVHGYRPSQHIISIEGHTEGPFKLTGAICYDATDIKLAADLRDLTDMFVIAAYNKDVDTFDNMASALQ<br>WHMYQHVIITNTGEYGGSTMQAPYKEKHHKLISHAHGTGQIAISTADIDLAAFRRKIKEYKKTKTQPAGFKRKH |
| DRT1<br>R582D,<br>K608E | MKLLNKKYYNLEPNFDYLKDSFILGLAWKKTDRFVRTHNWYADLLALDKCAFDISDEVTSWSSEAVNRNLSKSDIELIPAPKRASWFFNKGKWTTNKDDRKLRPLANISIKDQS<br>FATAVTMCLADAIETRQKDCSLRNLGYAEHVNKKVVSYGNRLVCDWDNERARFRWGGSEYYRKFSTDYRNFLQRPVYIGRETVNKISEIDDVYIISLDLKNFFGSIKISTLLEKLK<br>KISSDHYDHNIINENEFWGLASQILNWEWPEDSLSLLKNLDIEEENVGLPQGLASAGALANAYLIEFDESLTSNLRTKIDGSQIVLHDYCRYVDDIRLVISGEALTNEEIKKSIHEL<br>VQRILDETLDQDESDNEPYLKINDKKTYILGLSDIDNGSSLTNRINEIQNEVGTSSIPERNGLDNNIPALQQLLLTEQDNFLEDADGLFSGFNNDKSIKLESLRRFSAHRLETSLA<br>NKSKLISPAERKQFDNESALIAKKLLKTWLKDPSIMVIFRKAITINPNLDAYKTILEIIFLRIQHNREKCDRYIMLYLLSDIFRSVVDIYRKLSPECTSDYQKLMSEVTSFAHKLLSCDS<br>VIPNYAYQQALFYLAVINKPFIASKKSSSDLSKLQHVLIKRHLEPLSSSDGYLFELSAQISKDYQANVAFLLSHTNTTEVVDSIVEKFAYRGGEFWNSIWKEFVRTKDKVRINKFRW<br>AIPKNESKPNGSDHYLSSVISFKENPFKYEHALLKLGIALVNLLEDTEKNVWQSDGKQYSPHEIKVKLEGHSTSWTELWRQNSIISCSIDKIAQDPRYESPKWLVNCQQAKNDE<br>KKIYWVCSVLRSAALGNVDYTQRNDLKPDRVEYDGIHSQFYKRRMGMLHTPESIVGSYATITDWFASFLQHGLQWPGFSSSYINQEDILSITSLGEFKKCLLERLGYLNTQICTS<br>SSVPTLPTVVNRPELASNYFRIVTVQQLFPKDKHFHPSDVTLDNPEVRWKHREHLAEICKLTEQTLDAKLKTESRDHTSTADLIVFSELAVHPEDEDIVRALAFRTRSIIFCGFVFC<br>EQDGQIVNKARWIIPDSSESGTQWRVRDQGKFHMTSGEMSLGVHGYRPSQHIISIEGHTEGPFKLTGAICYDATDIKLAADLRDLTDMFVIAAYNKDVDTFDNMASALQWHM<br>YQHVIITNTGEYGGSTMQAPYKEKHHKLISHAHGTGQIAISTADIDLAAFRRKIKEYKKTKTQPAGFKRKH |
| DRT1<br>K791E | MKLLNKKYYNLEPNFDYLKDSFILGLAWKKTDRFVRTHNWYADLLALDKCAFDISDEVTSWSSEAVNRNLSKSDIELIPAPKRASWFFNKGKWTTNKDDRKLRPLANISIKDQS<br>FATAVTMCLADAIETRQKDCSLRNLGYAEHVNKKVVSYGNRLVCDWDNERARFRWGGSEYYRKFSTDYRNFLQRPVYIGRETVNKISEIDDVYIISLDLKNFFGSIKISTLLEKLK<br>KISSDHYDHNIINENEFWGLASQILNWEWPEDSLSLLKNLDIEEENVGLPQGLASAGALANAYLIEFDESLTSNLRTKIDGSQIVLHDYCRYVDDIRLVISGEALTNEEIKKSIHEL<br>VQRILDETLDQDESDNEPYLKINDKKTYILGLSDIDNGSSLTNRINEIQNEVGTSSIPERNGLDNNIPALQQLLLTEQDNFLEDADGLFSGFNNDKSIKLESLRRFSAHRLETSLA<br>NKSKLISPAERKQFDNESALIAKKLLKTWLKDPSIMVIFRKAITINPNLDAYKTILEIIFLRIQHNREKCDRYIMLYLLSDIFRSVVDIYRKLSPECTSDYQKLMSEVTSFAHKLLSCRS<br>VIPNYAYQQALFYLAVINKPFIASKKSSSDLSKLQHVLIKRHLEPLSSSDGYLFELSAQISKDYQANVAFLLSHTNTTEVVDSIVEKFAYRGGEFWNSIWKEFVRTKDKVRINKFRW<br>AIPKNESKPNGSDHYLSSVISFKENPFKYEHALLKLGIALVNLLEDTEKNVWQSDGKQYSPHEIKVKLEGHSTSWTELWRQNSIISCSIDEIAQDPRYESPKWLVNCQQAKNDE<br>KKIYWVCSVLRSAALGNVDYTQRNDLKPDRVEYDGIHSQFYKRRMGMLHTPESIVGSYATITDWFASFLQHGLQWPGFSSSYINQEDILSITSLGEFKKCLLERLGYLNTQICTS<br>SSVPTLPTVVNRPELASNYFRIVTVQQLFPKDKHFHPSDVTLDNPEVRWKHREHLAEICKLTEQTLDAKLKTESRDHTSTADLIVFSELAVHPEDEDIVRALAFRTRSIIFCGFVFC<br>EQDGQIVNKARWIIPDSSESGTQWRVRDQGKFHMTSGEMSLGVHGYRPSQHIISIEGHTEGPFKLTGAICYDATDIKLAADLRDLTDMFVIAAYNKDVDTFDNMASALQWHM<br>YQHVIITNTGEYGGSTMQAPYKEKHHKLISHAHGTGQIAISTADIDLAAFRRKIKEYKKTKTQPAGFKRKH |
| DRT1<br>E1214K,<br>I1212A | MKLLNKKYYNLEPNFDYLKDSFILGLAWKKTDRFVRTHNWYADLLALDKCAFDISDEVTSWSSEAVNRNLSKSDIELIPAPKRASWFFNKGKWTTNKDDRKLRPLANISIKDQS<br>FATAVTMCLADAIETRQKDCSLRNLGYAEHVNKKVVSYGNRLVCDWDNERARFRWGGSEYYRKFSTDYRNFLQRPVYIGRETVNKISEIDDVYIISLDLKNFFGSIKISTLLEKLK<br>KISSDHYDHNIINENEFWGLASQILNWEWPEDSLSLLKNLDIEEENVGLPQGLASAGALANAYLIEFDESLTSNLRTKIDGSQIVLHDYCRYVDDIRLVISGEALTNEEIKKSIHEL<br>VQRILDETLDQDESDNEPYLKINDKKTYILGLSDIDNGSSLTNRINEIQNEVGTSSIPERNGLDNNIPALQQLLLTEQDNFLEDADGLFSGFNNDKSIKLESLRRFSAHRLETSLA<br>NKSKLISPAERKQFDNESALIAKKLLKTWLKDPSIMVIFRKAITINPNLDAYKTILEIIFLRIQHNREKCDRYIMLYLLSDIFRSVVDIYRKLSPECTSDYQKLMSEVTSFAHKLLSCRS<br>VIPNYAYQQALFYLAVINKPFIASKKSSSDLSKLQHVLIKRHLEPLSSSDGYLFELSAQISKDYQANVAFLLSHTNTTEVVDSIVEKFAYRGGEFWNSIWKEFVRTKDKVRINKFRW<br>AIPKNESKPNGSDHYLSSVISFKENPFKYEHALLKLGIALVNLLEDTEKNVWQSDGKQYSPHEIKVKLEGHSTSWTELWRQNSIISCSIDKIAQDPRYESPKWLVNCQQAKNDE<br>KKIYWVCSVLRSAALGNVDYTQRNDLKPDRVEYDGIHSQFYKRRMGMLHTPESIVGSYATITDWFASFLQHGLQWPGFSSSYINQEDILSITSLGEFKKCLLERLGYLNTQICTS<br>SSVPTLPTVVNRPELASNYFRIVTVQQLFPKDKHFHPSDVTLDNPEVRWKHREHLAEICKLTEQTLDAKLKTESRDHTSTADLIVFSELAVHPEDEDIVRALAFRTRSIIFCGFVFC<br>EQDGQIVNKARWIIPDSSESGTQWRVRDQGKFHMTSGEMSLGVHGYRPSQHIISIEGHTEGPFKLTGAICYDATDIKLAADLRDLTDMFVIAAYNKDVDTFDNMASALQWHM<br>YQHVIITNTGEYGGSTMQAPYKEKHHKLISHAHGTGQIAISTADIDLAAFRRKAKKYKKTKTQPAGFKRKH |
| DRT1<br>K1213E | MKLLNKKYYNLEPNFDYLKDSFILGLAWKKTDRFVRTHNWYADLLALDKCAFDISDEVTSWSSEAVNRNLSKSDIELIPAPKRASWFFNKGKWTTNKDDRKLRPLANISIKDQS<br>FATAVTMCLADAIETRQKDCSLRNLGYAEHVNKKVVSYGNRLVCDWDNERARFRWGGSEYYRKFSTDYRNFLQRPVYIGRETVNKISEIDDVYIISLDLKNFFGSIKISTLLEKLK<br>KISSDHYDHNIINENEFWGLASQILNWEWPEDSLSLLKNLDIEEENVGLPQGLASAGALANAYLIEFDESLTSNLRTKIDGSQIVLHDYCRYVDDIRLVISGEALTNEEIKKSIHEL<br>VQRILDETLDQDESDNEPYLKINDKKTYILGLSDIDNGSSLTNRINEIQNEVGTSSIPERNGLDNNIPALQQLLLTEQDNFLEDADGLFSGFNNDKSIKLESLRRFSAHRLETSLA<br>NKSKLISPAERKQFDNESALIAKKLLKTWLKDPSIMVIFRKAITINPNLDAYKTILEIIFLRIQHNREKCDRYIMLYLLSDIFRSVVDIYRKLSPECTSDYQKLMSEVTSFAHKLLSCRS<br>VIPNYAYQQALFYLAVINKPFIASKKSSSDLSKLQHVLIKRHLEPLSSSDGYLFELSAQISKDYQANVAFLLSHTNTTEVVDSIVEKFAYRGGEFWNSIWKEFVRTKDKVRINKFRW<br>AIPKNESKPNGSDHYLSSVISFKENPFKYEHALLKLGIALVNLLEDTEKNVWQSDGKQYSPHEIKVKLEGHSTSWTELWRQNSIISCSIDKIAQDPRYESPKWLVNCQQAKNDE<br>KKIYWVCSVLRSAALGNVDYTQRNDLKPDRVEYDGIHSQFYKRRMGMLHTPESIVGSYATITDWFASFLQHGLQWPGFSSSYINQEDILSITSLGEFKKCLLERLGYLNTQICTS<br>SSVPTLPTVVNRPELASNYFRIVTVQQLFPKDKHFHPSDVTLDNPEVRWKHREHLAEICKLTEQTLDAKLKTESRDHTSTADLIVFSELAVHPEDEDIVRALAFRTRSIIFCGFVFC<br>EQDGQIVNKARWIIPDSSESGTQWRVRDQGKFHMTSGEMSLGVHGYRPSQHIISIEGHTEGPFKLTGAICYDATDIKLAADLRDLTDMFVIAAYNKDVDTFDNMASALQWHM<br>YQHVIITNTGEYGGSTMQAPYKEKHHKLISHAHGTGQIAISTADIDLAAFRRKIEEYKKTKTQPAGFKRKH |
| DRT1<br>K1213E,<br>E1214K | MKLLNKKYYNLEPNFDYLKDSFILGLAWKKTDRFVRTHNWYADLLALDKCAFDISDEVTSWSSEAVNRNLSKSDIELIPAPKRASWFFNKGKWTTNKDDRKLRPLANISIKDQS<br>FATAVTMCLADAIETRQKDCSLRNLGYAEHVNKKVVSYGNRLVCDWDNERARFRWGGSEYYRKFSTDYRNFLQRPVYIGRETVNKISEIDDVYIISLDLKNFFGSIKISTLLEKLK<br>KISSDHYDHNIINENEFWGLASQILNWEWPEDSLSLLKNLDIEEENVGLPQGLASAGALANAYLIEFDESLTSNLRTKIDGSQIVLHDYCRYVDDIRLVISGEALTNEEIKKSIHEL<br>VQRILDETLDQDESDNEPYLKINDKKTYILGLSDIDNGSSLTNRINEIQNEVGTSSIPERNGLDNNIPALQQLLLTEQDNFLEDADGLFSGFNNDKSIKLESLRRFSAHRLETSLA<br>NKSKLISPAERKQFDNESALIAKKLLKTWLKDPSIMVIFRKAITINPNLDAYKTILEIIFLRIQHNREKCDRYIMLYLLSDIFRSVVDIYRKLSPECTSDYQKLMSEVTSFAHKLLSCRS<br>VIPNYAYQQALFYLAVINKPFIASKKSSSDLSKLQHVLIKRHLEPLSSSDGYLFELSAQISKDYQANVAFLLSHTNTTEVVDSIVEKFAYRGGEFWNSIWKEFVRTKDKVRINKFRW |

| | |
|---|---|
| | AIPKNESKPNGSDHYLSSVISFKENPFKYEHALLKLGIALVNLLEDTEKNVWQSDGKQYSPHEIKVKLEGHSTSWTELWRQNSIISCSIDKIAQDPRYESPKWLVNCQQAKNDE<br>KKIYWVCSVLRSAALGNVDYTQRNDLKPDRVEYDGIHSQFYKRRMGMLHTPESIVGSYATITDWFASFLQHGLQWPGFSSSYINQEDILSITSLGEFKKCLLERLGYLNTQICTS<br>SSVPTLPTVVNRPELASNYFRIVTVQQLFPKDKHFHPSDVTLDNPEVRWKHREHLAEICKLTEQTLDAKLKTESRDHTSTADLIVFSELAVHPEDEDIVRALAFRTRSIIFCGFVFC<br>EQDGQIVNKARWIIPDSSESGTQWRVRDQGKFHMTSGEMSLGVHGYRPSQHIISIEGHTEGPFKLTGAICYDATDIKLAADLRDLTDMFVIAAYNKDVDTFDNMASALQWHM<br>YQHVIITNTGEYGGSTMQAPYKEKHHKLISHAHGTGQIAISTADIDLAAFRRKIEKYKKTKTQPAGFKRKH |
| DRT1<br>E1214K | MKLLNKKYYNLEPNFDYLKDSFILGLAWKKTDRFVRTHNWYADLLALDKCAFDISDEVTSWSSEAVNRNLSKSDIELIPAPKRASWFFNKGKWTTNKDDRKLRPLANISIKDQS<br>FATAVTMCLADAIETRQKDCSLRNLGYAEHVNKKVVSYGNRLVCDWDNERARFRWGGSEYYRKFSTDYRNFLQRPVYIGRETVNKISEIDDVYIISLDLKNFFGSIKISTLLEKLK<br>KISSDHYDHNIINENEFWGLASQILNWEWPEDSLSLLKNLDIEEENVGLPQGLASAGALANAYLIEFDESLTSNLRTKIDGSQIVLHDYCRYVDDIRLVISGEALTNEEIKKSIHEL<br>VQRILDETLDQDESDNEPYLKINDKKTYILGLSDIDNGSSLTNRINEIQNEVGTSSIPERNGLDNNIPALQQLLLTEQDNFLEDADGLFSGFNNDKSIKLESLRRFSAHRLETSLA<br>NKSKLISPAERKQFDNESALIAKKLLKTWLKDPSIMVIFRKAITINPNLDAYKTILEIIFLRIQHNREKCDRYIMLYLLSDIFRSVVDIYRKLSPECTSDYQKLMSEVTSFAHKLLSCRS<br>VIPNYAYQQALFYLAVINKPFIASKKSSSDLSKLQHVLIKRHLEPLSSSDGYLFELSAQISKDYQANVAFLLSHTNTTEVVDSIVEKFAYRGGEFWNSIWKEFVRTKDKVRINKFRW<br>AIPKNESKPNGSDHYLSSVISFKENPFKYEHALLKLGIALVNLLEDTEKNVWQSDGKQYSPHEIKVKLEGHSTSWTELWRQNSIISCSIDKIAQDPRYESPKWLVNCQQAKNDE<br>KKIYWVCSVLRSAALGNVDYTQRNDLKPDRVEYDGIHSQFYKRRMGMLHTPESIVGSYATITDWFASFLQHGLQWPGFSSSYINQEDILSITSLGEFKKCLLERLGYLNTQICTS<br>SSVPTLPTVVNRPELASNYFRIVTVQQLFPKDKHFHPSDVTLDNPEVRWKHREHLAEICKLTEQTLDAKLKTESRDHTSTADLIVFSELAVHPED EDIVRALAFRTRSIIFCGFVFC<br>EQDGQIVNKARWIIPDSSESGTQWRVRDQGKFHMTSGEMSLGVHGYRPSQHIISIEGHTEGPFKLTGAICYDATDIKLAADLRDLTDMFVIAAYNKDVDTFDNMASALQWHM<br>YQHVIITNTGEYGGSTMQAPYKEKHHKLISHAHGTGQIAISTADIDLAAFRRKIKKYKKTKTQPAGFKRKH |
| DRT1<br>K1217E | MKLLNKKYYNLEPNFDYLKDSFILGLAWKKTDRFVRTHNWYADLLALDKCAFDISDEVTSWSSEAVNRNLSKSDIELIPAPKRASWFFNKGKWTTNKDDRKLRPLANISIKDQS<br>FATAVTMCLADAIETRQKDCSLRNLGYAEHVNKKVVSYGNRLVCDWDNERARFRWGGSEYYRKFSTDYRNFLQRPVYIGRETVNKISEIDDVYIISLDLKNFFGSIKISTLLEKLK<br>KISSDHYDHNIINENEFWGLASQILNWEWPEDSLSLLKNLDIEEENVGLPQGLASAGALANAYLIEFDESLTSNLRTKIDGSQIVLHDYCRYVDDIRLVISGEALTNEEIKKSIHEL<br>VQRILDETLDQDESDNEPYLKINDKKTYILGLSDIDNGSSLTNRINEIQNEVGTSSIPERNGLDNNIPALQQLLLTEQDNFLEDADGLFSGFNNDKSIKLESLRRFSAHRLETSLA<br>NKSKLISPAERKQFDNESALIAKKLLKTWLKDPSIMVIFRKAITINPNLDAYKTILEIIFLRIQHNREKCDRYIMLYLLSDIFRSVVDIYRKLSPECTSDYQKLMSEVTSFAHKLLSCRS<br>VIPNYAYQQALFYLAVINKPFIASKKSSSDLSKLQHVLIKRHLEPLSSSDGYLFELSAQISKDYQANVAFLLSHTNTTEVVDSIVEKFAYRGGEFWNSIWKEFVRTKDKVRINKFRW<br>AIPKNESKPNGSDHYLSSVISFKENPFKYEHALLKLGIALVNLLEDTEKNVWQSDGKQYSPHEIKVKLEGHSTSWTELWRQNSIISCSIDKIAQDPRYESPKWLVNCQQAKNDE<br>KKIYWVCSVLRSAALGNVDYTQRNDLKPDRVEYDGIHSQFYKRRMGMLHTPESIVGSYATITDWFASFLQHGLQWPGFSSSYINQEDILSITSLGEFKKCLLERLGYLNTQICTS<br>SSVPTLPTVVNRPELASNYFRIVTVQQLFPKDKHFHPSDVTLDNPEVRWKHREHLAEICKLTEQTLDAKLKTESRDHTSTADLIVFSELAVHPED EDIVRALAFRTRSIIFCGFVFC<br>EQDGQIVNKARWIIPDSSESGTQWRVRDQGKFHMTSGEMSLGVHGYRPSQHIISIEGHTEGPFKLTGAICYDATDIKLAADLRDLTDMFVIAAYNKDVDTFDNMASALQWHM<br>YQHVIITNTGEYGGSTMQAPYKEKHHKLISHAHGTGQIAISTADIDLAAFRRKIKEYKETKTQPAGFKRKH |
| DRT1<br>K1219A | MKLLNKKYYNLEPNFDYLKDSFILGLAWKKTDRFVRTHNWYADLLALDKCAFDISDEVTSWSSEAVNRNLSKSDIELIPAPKRASWFFNKGKWTTNKDDRKLRPLANISIKDQS<br>FATAVTMCLADAIETRQKDCSLRNLGYAEHVNKKVVSYGNRLVCDWDNERARFRWGGSEYYRKFSTDYRNFLQRPVYIGRETVNKISEIDDVYIISLDLKNFFGSIKISTLLEKLK<br>KISSDHYDHNIINENEFWGLASQILNWEWPEDSLSLLKNLDIEEENVGLPQGLASAGALANAYLIEFDESLTSNLRTKIDGSQIVLHDYCRYVDDIRLVISGEALTNEEIKKSIHEL<br>VQRILDETLDQDESDNEPYLKINDKKTYILGLSDIDNGSSLTNRINEIQNEVGTSSIPERNGLDNNIPALQQLLLTEQDNFLEDADGLFSGFNNDKSIKLESLRRFSAHRLETSLA<br>NKSKLISPAERKQFDNESALIAKKLLKTWLKDPSIMVIFRKAITINPNLDAYKTILEIIFLRIQHNREKCDRYIMLYLLSDIFRSVVDIYRKLSPECTSDYQKLMSEVTSFAHKLLSCRS<br>VIPNYAYQQALFYLAVINKPFIASKKSSSDLSKLQHVLIKRHLEPLSSSDGYLFELSAQISKDYQANVAFLLSHTNTTEVVDSIVEKFAYRGGEFWNSIWKEFVRTKDKVRINKFRW<br>AIPKNESKPNGSDHYLSSVISFKENPFKYEHALLKLGIALVNLLEDTEKNVWQSDGKQYSPHEIKVKLEGHSTSWTELWRQNSIISCSIDKIAQDPRYESPKWLVNCQQAKNDE<br>KKIYWVCSVLRSAALGNVDYTQRNDLKPDRVEYDGIHSQFYKRRMGMLHTPESIVGSYATITDWFASFLQHGLQWPGFSSSYINQEDILSITSLGEFKKCLLERLGYLNTQICTS<br>SSVPTLPTVVNRPELASNYFRIVTVQQLFPKDKHFHPSDVTLDNPEVRWKHREHLAEICKLTEQTLDAKLKTESRDHTSTADLIVFSELAVHPED EDIVRALAFRTRSIIFCGFVFC<br>EQDGQIVNKARWIIPDSSESGTQWRVRDQGKFHMTSGEMSLGVHGYRPSQHIISIEGHTEGPFKLTGAICYDATDIKLAADLRDLTDMFVIAAYNKDVDTFDNMASALQWHM<br>YQHVIITNTGEYGGSTMQAPYKEKHHKLISHAHGTGQIAISTADIDLAAFRRKIKEYKKTATQPAGFKRKH |
| DRT1<br>A1223F | MKLLNKKYYNLEPNFDYLKDSFILGLAWKKTDRFVRTHNWYADLLALDKCAFDISDEVTSWSSEAVNRNLSKSDIELIPAPKRASWFFNKGKWTTNKDDRKLRPLANISIKDQS<br>FATAVTMCLADAIETRQKDCSLRNLGYAEHVNKKVVSYGNRLVCDWDNERARFRWGGSEYYRKFSTDYRNFLQRPVYIGRETVNKISEIDDVYIISLDLKNFFGSIKISTLLEKLK<br>KISSDHYDHNIINENEFWGLASQILNWEWPEDSLSLLKNLDIEEENVGLPQGLASAGALANAYLIEFDESLTSNLRTKIDGSQIVLHDYCRYVDDIRLVISGEALTNEEIKKSIHEL<br>VQRILDETLDQDESDNEPYLKINDKKTYILGLSDIDNGSSLTNRINEIQNEVGTSSIPERNGLDNNIPALQQLLLTEQDNFLEDADGLFSGFNNDKSIKLESLRRFSAHRLETSLA<br>NKSKLISPAERKQFDNESALIAKKLLKTWLKDPSIMVIFRKAITINPNLDAYKTILEIIFLRIQHNREKCDRYIMLYLLSDIFRSVVDIYRKLSPECTSDYQKLMSEVTSFAHKLLSCRS<br>VIPNYAYQQALFYLAVINKPFIASKKSSSDLSKLQHVLIKRHLEPLSSSDGYLFELSAQISKDYQANVAFLLSHTNTTEVVDSIVEKFAYRGGEFWNSIWKEFVRTKDKVRINKFRW<br>AIPKNESKPNGSDHYLSSVISFKENPFKYEHALLKLGIALVNLLEDTEKNVWQSDGKQYSPHEIKVKLEGHSTSWTELWRQNSIISCSIDKIAQDPRYESPKWLVNCQQAKNDE<br>KKIYWVCSVLRSAALGNVDYTQRNDLKPDRVEYDGIHSQFYKRRMGMLHTPESIVGSYATITDWFASFLQHGLQWPGFSSSYINQEDILSITSLGEFKKCLLERLGYLNTQICTS<br>SSVPTLPTVVNRPELASNYFRIVTVQQLFPKDKHFHPSDVTLDNPEVRWKHREHLAEICKLTEQTLDAKLKTESRDHTSTADLIVFSELAVHPED EDIVRALAFRTRSIIFCGFVFC<br>EQDGQIVNKARWIIPDSSESGTQWRVRDQGKFHMTSGEMSLGVHGYRPSQHIISIEGHTEGPFKLTGAICYDATDIKLAADLRDLTDMFVIAAYNKDVDTFDNMASALQWHM<br>YQHVIITNTGEYGGSTMQAPYKEKHHKLISHAHGTGQIAISTADIDLAAFRRKIKEYKKTKTQPFGFKRKH |
| DRT1<br>R1227A | MKLLNKKYYNLEPNFDYLKDSFILGLAWKKTDRFVRTHNWYADLLALDKCAFDISDEVTSWSSEAVNRNLSKSDIELIPAPKRASWFFNKGKWTTNKDDRKLRPLANISIKDQS<br>FATAVTMCLADAIETRQKDCSLRNLGYAEHVNKKVVSYGNRLVCDWDNERARFRWGGSEYYRKFSTDYRNFLQRPVYIGRETVNKISEIDDVYIISLDLKNFFGSIKISTLLEKLK<br>KISSDHYDHNIINENEFWGLASQILNWEWPEDSLSLLKNLDIEEENVGLPQGLASAGALANAYLIEFDESLTSNLRTKIDGSQIVLHDYCRYVDDIRLVISGEALTNEEIKKSIHEL<br>VQRILDETLDQDESDNEPYLKINDKKTYILGLSDIDNGSSLTNRINEIQNEVGTSSIPERNGLDNNIPALQQLLLTEQDNFLEDADGLFSGFNNDKSIKLESLRRFSAHRLETSLA<br>NKSKLISPAERKQFDNESALIAKKLLKTWLKDPSIMVIFRKAITINPNLDAYKTILEIIFLRIQHNREKCDRYIMLYLLSDIFRSVVDIYRKLSPECTSDYQKLMSEVTSFAHKLLSCRS<br>VIPNYAYQQALFYLAVINKPFIASKKSSSDLSKLQHVLIKRHLEPLSSSDGYLFELSAQISKDYQANVAFLLSHTNTTEVVDSIVEKFAYRGGEFWNSIWKEFVRTKDKVRINKFRW<br>AIPKNESKPNGSDHYLSSVISFKENPFKYEHALLKLGIALVNLLEDTEKNVWQSDGKQYSPHEIKVKLEGHSTSWTELWRQNSIISCSIDKIAQDPRYESPKWLVNCQQAKNDE<br>KKIYWVCSVLRSAALGNVDYTQRNDLKPDRVEYDGIHSQFYKRRMGMLHTPESIVGSYATITDWFASFLQHGLQWPGFSSSYINQEDILSITSLGEFKKCLLERLGYLNTQICTS<br>SSVPTLPTVVNRPELASNYFRIVTVQQLFPKDKHFHPSDVTLDNPEVRWKHREHLAEICKLTEQTLDAKLKTESRDHTSTADLIVFSELAVHPED EDIVRALAFRTRSIIFCGFVFC<br>EQDGQIVNKARWIIPDSSESGTQWRVRDQGKFHMTSGEMSLGVHGYRPSQHIISIEGHTEGPFKLTGAICYDATDIKLAADLRDLTDMFVIAAYNKDVDTFDNMASALQWHM<br>YQHVIITNTGEYGGSTMQAPYKEKHHKLISHAHGTGQIAISTADIDLAAFRRKIKEYKKTKTQPAGFKAKH |
| DRT1 ΔC8 | MKLLNKKYYNLEPNFDYLKDSFILGLAWKKTDRFVRTHNWYADLLALDKCAFDISDEVTSWSSEAVNRNLSKSDIELIPAPKRASWFFNKGKWTTNKDDRKLRPLANISIKDQS<br>FATAVTMCLADAIETRQKDCSLRNLGYAEHVNKKVVSYGNRLVCDWDNERARFRWGGSEYYRKFSTDYRNFLQRPVYIGRETVNKISEIDDVYIISLDLKNFFGSIKISTLLEKLK<br>KISSDHYDHNIINENEFWGLASQILNWEWPEDSLSLLKNLDIEEENVGLPQGLASAGALANAYLIEFDESLTSNLRTKIDGSQIVLHDYCRYVDDIRLVISGEALTNEEIKKSIHEL<br>VQRILDETLDQDESDNEPYLKINDKKTYILGLSDIDNGSSLTNRINEIQNEVGTSSIPERNGLDNNIPALQQLLLTEQDNFLEDADGLFSGFNNDKSIKLESLRRFSAHRLETSLA<br>NKSKLISPAERKQFDNESALIAKKLLKTWLKDPSIMVIFRKAITINPNLDAYKTILEIIFLRIQHNREKCDRYIMLYLLSDIFRSVVDIYRKLSPECTSDYQKLMSEVTSFAHKLLSCRS<br>VIPNYAYQQALFYLAVINKPFIASKKSSSDLSKLQHVLIKRHLEPLSSSDGYLFELSAQISKDYQANVAFLLSHTNTTEVVDSIVEKFAYRGGEFWNSIWKEFVRTKDKVRINKFRW<br>AIPKNESKPNGSDHYLSSVISFKENPFKYEHALLKLGIALVNLLEDTEKNVWQSDGKQYSPHEIKVKLEGHSTSWTELWRQNSIISCSIDKIAQDPRYESPKWLVNCQQAKNDE<br>KKIYWVCSVLRSAALGNVDYTQRNDLKPDRVEYDGIHSQFYKRRMGMLHTPESIVGSYATITDWFASFLQHGLQWPGFSSSYINQEDILSITSLGEFKKCLLERLGYLNTQICTS<br>SSVPTLPTVVNRPELASNYFRIVTVQQLFPKDKHFHPSDVTLDNPEVRWKHREHLAEICKLTEQTLDAKLKTESRDHTSTADLIVFSELAVHPED EDIVRALAFRTRSIIFCGFVFC<br>EQDGQIVNKARWIIPDSSESGTQWRVRDQGKFHMTSGEMSLGVHGYRPSQHIISIEGHTEGPFKLTGAICYDATDIKLAADLRDLTDMFVIAAYNKDVDTFDNMASALQWHM<br>YQHVIITNTGEYGGSTMQAPYKEKHHKLISHAHGTGQIAISTADIDLAAFRRKIKEYKKTKTQ |
| DRT1<br>ΔC12 | MKLLNKKYYNLEPNFDYLKDSFILGLAWKKTDRFVRTHNWYADLLALDKCAFDISDEVTSWSSEAVNRNLSKSDIELIPAPKRASWFFNKGKWTTNKDDRKLRPLANISIKDQS<br>FATAVTMCLADAIETRQKDCSLRNLGYAEHVNKKVVSYGNRLVCDWDNERARFRWGGSEYYRKFSTDYRNFLQRPVYIGRETVNKISEIDDVYIISLDLKNFFGSIKISTLLEKLK<br>KISSDHYDHNIINENEFWGLASQILNWEWPEDSLSLLKNLDIEEENVGLPQGLASAGALANAYLIEFDESLTSNLRTKIDGSQIVLHDYCRYVDDIRLVISGEALTNEEIKKSIHEL<br>VQRILDETLDQDESDNEPYLKINDKKTYILGLSDIDNGSSLTNRINEIQNEVGTSSIPERNGLDNNIPALQQLLLTEQDNFLEDADGLFSGFNNDKSIKLESLRRFSAHRLETSLA<br>NKSKLISPAERKQFDNESALIAKKLLKTWLKDPSIMVIFRKAITINPNLDAYKTILEIIFLRIQHNREKCDRYIMLYLLSDIFRSVVDIYRKLSPECTSDYQKLMSEVTSFAHKLLSCRS<br>VIPNYAYQQALFYLAVINKPFIASKKSSSDLSKLQHVLIKRHLEPLSSSDGYLFELSAQISKDYQANVAFLLSHTNTTEVVDSIVEKFAYRGGEFWNSIWKEFVRTKDKVRINKFRW<br>AIPKNESKPNGSDHYLSSVISFKENPFKYEHALLKLGIALVNLLEDTEKNVWQSDGKQYSPHEIKVKLEGHSTSWTELWRQNSIISCSIDKIAQDPRYESPKWLVNCQQAKNDE<br>KKIYWVCSVLRSAALGNVDYTQRNDLKPDRVEYDGIHSQFYKRRMGMLHTPESIVGSYATITDWFASFLQHGLQWPGFSSSYINQEDILSITSLGEFKKCLLERLGYLNTQICTS<br>SSVPTLPTVVNRPELASNYFRIVTVQQLFPKDKHFHPSDVTLDNPEVRWKHREHLAEICKLTEQTLDAKLKTESRDHTSTADLIVFSELAVHPED EDIVRALAFRTRSIIFCGFVFC<br>EQDGQIVNKARWIIPDSSESGTQWRVRDQGKFHMTSGEMSLGVHGYRPSQHIISIEGHTEGPFKLTGAICYDATDIKLAADLRDLTDMFVIAAYNKDVDTFDNMASALQWHM<br>YQHVIITNTGEYGGSTMQAPYKEKHHKLISHAHGTGQIAISTADIDLAAFRRKIKEYKK |
| DRT1<br>I200A,<br>I792A | MKLLNKKYYNLEPNFDYLKDSFILGLAWKKTDRFVRTHNWYADLLALDKCAFDISDEVTSWSSEAVNRNLSKSDIELIPAPKRASWFFNKGKWTTNKDDRKLRPLANISIKDQS<br>FATAVTMCLADAIETRQKDCSLRNLGYAEHVNKKVVSYGNRLVCDWDNERARFRWGGSEYYRKFSTDYRNFLQRPVYIGRETVNKASEIDDVYIISLDLKNFFGSIKISTLLEKL<br>KKISSDHYDHNIINENEFWGLASQILNWEWPEDSLSLLKNLDIEEENVGLPQGLASAGALANAYLIEFDESLTSNLRTKIDGSQIVLHDYCRYVDDIRLVISGEALTNEEIKKSIHE<br>LVQRILDETLDQDESDNEPYLKINDKKTYILGLSDIDNGSSLTNRINEIQNEVGTSSIPERNGLDNNIPALQQLLLTEQDNFLEDADGLFSGFNNDKSIKLESLRRFSAHRLETSL<br>ANKSKLISPAERKQFDNESALIAKKLLKTWLKDPSIMVIFRKAITINPNLDAYKTILEIIFLRIQHNREKCDRYIMLYLLSDIFRSVVDIYRKLSPECTSDYQKLMSEVTSFAHKLLSCR |

| | |
|---|---|
| | SVIPNYAYQQALFYLAVINKPFIASKKSSSDLSKLQHVLIKRHLEPLSSSDGYLFELSAQISKDYQANVAFLLSHTNTTEVVDSIVEKFAYRGGEFWNSIWKEFVRTKDKVRINKFR<br>WAIPKNESKPNGSDHYLSSVISFKENPFKYEHALLKLGIALVNLLEDTEKNVWQSDGKQYSPHEIKVKLEGHSTSWTELWRQNSIISCSIDKAAQDPRYESPKWLVNCQQAKN<br>DEKKIYWVCSVLRSAALGNVDYTQRNDLKPDRVEYDGIHSQFYKRRMGMLHTPESIVGSYATITDWFASFLQHGLQWPGFSSSYINQEDILSITSLGEFKKCLLERLGYLNTQI<br>CTSSSVPTLPTVVNRPELASNYFRIVTVQQLFPKDKHFHPSDVTLDNPEVRWKHREHLAEICKLTEQTLDAKLKTESRDHTSTADLIVFSELAVHPEDEDIVRALAFRTRSIIFCGF<br>VFCEQDGQIVNKARWIIPDSSESGTQWRVRDQGKFHMTSGEMSLGVHGYRPSQHIISIEGHTEGPFKLTGAICYDATDIKLAADLRDLTDMFVIAAYNKDVDTFDNMASALQ<br>WHMYQHVIITNTGEYGGSTMQAPYKEKHHKLISHAHGTGQIAISTADIDLAAFRRKIKEYKKTKTQPAGFKRKH |
| DRT1<br>I309A | MKLLNKKYYNLEPNFDYLKDSFILGLAWKKTDRFVRTHNWYADLLALDKCAFDISDEVTSWSSEAVNRNLSKSDIELIPAPKRASWFFNKGKWTTNKDDRKLRPLANISIKDQS<br>FATAVTMCLADAIETRQKDCSLRNLGYAEHVNKKVVSYGNRLVCDWDNERARFRWGGSEYYRKFSTDYRNFLQRPVYIGRETVNKISEIDDVYIISLDLKNFFGSIKISTLLEKLK<br>KISSDHYDHNIINENEFWGLASQILNWEWPEDSLSLLKNLDIEEENVGLPQGLASAGALANAYLIEFDESLTSNLRTKADGSQIVLHDYCRYVDDIRLVISGEALTNEEIKKSIHE<br>LVQRILDETLDQDESDNEPYLKINDKKTYILGLSDIDNGSSLTNRINEIQNEVGTSSIPERNGLDNNIPALQQLLLTEQDNFLEDADGLFSGFNNDKSIKLESLRRFSAHRLETSL<br>ANKSKLISPAERKQFDNESALIAKKLLKTWLKDPSIMVIFRKAITINPNLDAYKTILEIIFLRIQHNREKCDRYIMLYLLSDIFRSVVDIYRKLSPECTSDYQKLMSEVTSFAHKLLSCR<br>SVIPNYAYQQALFYLAVINKPFIASKKSSSDLSKLQHVLIKRHLEPLSSSDGYLFELSAQISKDYQANVAFLLSHTNTTEVVDSIVEKFAYRGGEFWNSIWKEFVRTKDKVRINKFR<br>WAIPKNESKPNGSDHYLSSVISFKENPFKYEHALLKLGIALVNLLEDTEKNVWQSDGKQYSPHEIKVKLEGHSTSWTELWRQNSIISCSIDKIAQDPRYESPKWLVNCQQAKN<br>DEKKIYWVCSVLRSAALGNVDYTQRNDLKPDRVEYDGIHSQFYKRRMGMLHTPESIVGSYATITDWFASFLQHGLQWPGFSSSYINQEDILSITSLGEFKKCLLERLGYLNTQI<br>CTSSSVPTLPTVVNRPELASNYFRIVTVQQLFPKDKHFHPSDVTLDNPEVRWKHREHLAEICKLTEQTLDAKLKTESRDHTSTADLIVFSELAVHPEDEDIVRALAFRTRSIIFCGF<br>VFCEQDGQIVNKARWIIPDSSESGTQWRVRDQGKFHMTSGEMSLGVHGYRPSQHIISIEGHTEGPFKLTGAICYDATDIKLAADLRDLTDMFVIAAYNKDVDTFDNMASALQ<br>WHMYQHVIITNTGEYGGSTMQAPYKEKHHKLISHAHGTGQIAISTADIDLAAFRRKIKEYKKTKTQPAGFKRKH |
| DRT1<br>E333R | MKLLNKKYYNLEPNFDYLKDSFILGLAWKKTDRFVRTHNWYADLLALDKCAFDISDEVTSWSSEAVNRNLSKSDIELIPAPKRASWFFNKGKWTTNKDDRKLRPLANISIKDQS<br>FATAVTMCLADAIETRQKDCSLRNLGYAEHVNKKVVSYGNRLVCDWDNERARFRWGGSEYYRKFSTDYRNFLQRPVYIGRETVNKISEIDDVYIISLDLKNFFGSIKISTLLEKLK<br>KISSDHYDHNIINENEFWGLASQILNWEWPEDSLSLLKNLDIEEENVGLPQGLASAGALANAYLIEFDESLTSNLRTKIDGSQIVLHDYCRYVDDIRLVISGRALTNEEIKKSIHEL<br>VQRILDETLDQDESDNEPYLKINDKKTYILGLSDIDNGSSLTNRINEIQNEVGTSSIPERNGLDNNIPALQQLLLTEQDNFLEDADGLFSGFNNDKSIKLESLRRFSAHRLETSLA<br>NKSKLISPAERKQFDNESALIAKKLLKTWLKDPSIMVIFRKAITINPNLDAYKTILEIIFLRIQHNREKCDRYIMLYLLSDIFRSVVDIYRKLSPECTSDYQKLMSEVTSFAHKLLSCRS<br>VIPNYAYQQALFYLAVINKPFIASKKSSSDLSKLQHVLIKRHLEPLSSSDGYLFELSAQISKDYQANVAFLLSHTNTTEVVDSIVEKFAYRGGEFWNSIWKEFVRTKDKVRINKFRW<br>AIPKNESKPNGSDHYLSSVISFKENPFKYEHALLKLGIALVNLLEDTEKNVWQSDGKQYSPHEIKVKLEGHSTSWTELWRQNSIISCSIDKIAQDPRYESPKWLVNCQQAKNDE<br>KKIYWVCSVLRSAALGNVDYTQRNDLKPDRVEYDGIHSQFYKRRMGMLHTPESIVGSYATITDWFASFLQHGLQWPGFSSSYINQEDILSITSLGEFKKCLLERLGYLNTQICTS<br>SSVPTLPTVVNRPELASNYFRIVTVQQLFPKDKHFHPSDVTLDNPEVRWKHREHLAEICKLTEQTLDAKLKTESRDHTSTADLIVFSELAVHPEDEDIVRALAFRTRSIIFCGFVFC<br>EQDGQIVNKARWIIPDSSESGTQWRVRDQGKFHMTSGEMSLGVHGYRPSQHIISIEGHTEGPFKLTGAICYDATDIKLAADLRDLTDMFVIAAYNKDVDTFDNMASALQWHM<br>YQHVIITNTGEYGGSTMQAPYKEKHHKLISHAHGTGQIAISTADIDLAAFRRKIKEYKKTKTQPAGFKRKH |
| DRT1<br>R669A | MKLLNKKYYNLEPNFDYLKDSFILGLAWKKTDRFVRTHNWYADLLALDKCAFDISDEVTSWSSEAVNRNLSKSDIELIPAPKRASWFFNKGKWTTNKDDRKLRPLANISIKDQS<br>FATAVTMCLADAIETRQKDCSLRNLGYAEHVNKKVVSYGNRLVCDWDNERARFRWGGSEYYRKFSTDYRNFLQRPVYIGRETVNKISEIDDVYIISLDLKNFFGSIKISTLLEKLK<br>KISSDHYDHNIINENEFWGLASQILNWEWPEDSLSLLKNLDIEEENVGLPQGLASAGALANAYLIEFDESLTSNLRTKIDGSQIVLHDYCRYVDDIRLVISGEALTNEEIKKSIHEL<br>VQRILDETLDQDESDNEPYLKINDKKTYILGLSDIDNGSSLTNRINEIQNEVGTSSIPERNGLDNNIPALQQLLLTEQDNFLEDADGLFSGFNNDKSIKLESLRRFSAHRLETSLA<br>NKSKLISPAERKQFDNESALIAKKLLKTWLKDPSIMVIFRKAITINPNLDAYKTILEIIFLRIQHNREKCDRYIMLYLLSDIFRSVVDIYRKLSPECTSDYQKLMSEVTSFAHKLLSCRS<br>VIPNYAYQQALFYLAVINKPFIASKKSSSDLSKLQHVLIKRHLEPLSSSDGYLFELSAQISKDYQANVAFLLSHTNTTEVVDSIVEKFAYRGGEFWNSIWKEFVRTKDKVRINKFAW<br>AIPKNESKPNGSDHYLSSVISFKENPFKYEHALLKLGIALVNLLEDTEKNVWQSDGKQYSPHEIKVKLEGHSTSWTELWRQNSIISCSIDKIAQDPRYESPKWLVNCQQAKNDE<br>KKIYWVCSVLRSAALGNVDYTQRNDLKPDRVEYDGIHSQFYKRRMGMLHTPESIVGSYATITDWFASFLQHGLQWPGFSSSYINQEDILSITSLGEFKKCLLERLGYLNTQICTS<br>SSVPTLPTVVNRPELASNYFRIVTVQQLFPKDKHFHPSDVTLDNPEVRWKHREHLAEICKLTEQTLDAKLKTESRDHTSTADLIVFSELAVHPEDEDIVRALAFRTRSIIFCGFVFC<br>EQDGQIVNKARWIIPDSSESGTQWRVRDQGKFHMTSGEMSLGVHGYRPSQHIISIEGHTEGPFKLTGAICYDATDIKLAADLRDLTDMFVIAAYNKDVDTFDNMASALQWHM<br>YQHVIITNTGEYGGSTMQAPYKEKHHKLISHAHGTGQIAISTADIDLAAFRRKIKEYKKTKTQPAGFKRKH |
| DRT1<br>S967A | MKLLNKKYYNLEPNFDYLKDSFILGLAWKKTDRFVRTHNWYADLLALDKCAFDISDEVTSWSSEAVNRNLSKSDIELIPAPKRASWFFNKGKWTTNKDDRKLRPLANISIKDQS<br>FATAVTMCLADAIETRQKDCSLRNLGYAEHVNKKVVSYGNRLVCDWDNERARFRWGGSEYYRKFSTDYRNFLQRPVYIGRETVNKISEIDDVYIISLDLKNFFGSIKISTLLEKLK<br>KISSDHYDHNIINENEFWGLASQILNWEWPEDSLSLLKNLDIEEENVGLPQGLASAGALANAYLIEFDESLTSNLRTKIDGSQIVLHDYCRYVDDIRLVISGEALTNEEIKKSIHEL<br>VQRILDETLDQDESDNEPYLKINDKKTYILGLSDIDNGSSLTNRINEIQNEVGTSSIPERNGLDNNIPALQQLLLTEQDNFLEDADGLFSGFNNDKSIKLESLRRFSAHRLETSLA<br>NKSKLISPAERKQFDNESALIAKKLLKTWLKDPSIMVIFRKAITINPNLDAYKTILEIIFLRIQHNREKCDRYIMLYLLSDIFRSVVDIYRKLSPECTSDYQKLMSEVTSFAHKLLSCRS<br>VIPNYAYQQALFYLAVINKPFIASKKSSSDLSKLQHVLIKRHLEPLSSSDGYLFELSAQISKDYQANVAFLLSHTNTTEVVDSIVEKFAYRGGEFWNSIWKEFVRTKDKVRINKFRW<br>AIPKNESKPNGSDHYLSSVISFKENPFKYEHALLKLGIALVNLLEDTEKNVWQSDGKQYSPHEIKVKLEGHSTSWTELWRQNSIISCSIDKIAQDPRYESPKWLVNCQQAKNDE<br>KKIYWVCSVLRSAALGNVDYTQRNDLKPDRVEYDGIHSQFYKRRMGMLHTPESIVGSYATITDWFASFLQHGLQWPGFSSSYINQEDILSITSLGEFKKCLLERLGYLNTQICTS<br>SSVPTLPTVVNRPELASNYFRIVTVQQLFPKDKHFHPADVTLDNPEVRWKHREHLAEICKLTEQTLDAKLKTESRDHTSTADLIVFSELAVHPEDEDIVRALAFRTRSIIFCGFVFC<br>EQDGQIVNKARWIIPDSSESGTQWRVRDQGKFHMTSGEMSLGVHGYRPSQHIISIEGHTEGPFKLTGAICYDATDIKLAADLRDLTDMFVIAAYNKDVDTFDNMASALQWHM<br>YQHVIITNTGEYGGSTMQAPYKEKHHKLISHAHGTGQIAISTADIDLAAFRRKIKEYKKTKTQPAGFKRKH |
| DRT5L | MITKDIVVSAYNCLKSYAYYENLNFFLKEEIAQFEDSLYDEKIEKVVDFFENNNDINFNEWLNKIDVELLPKKIDSHLDIEQTSGALFLSNNKTSDEYKVGAVNYLIVAPIEIYLIETL<br>WSIFVGSMLDDKFSDSVYGNRISKVIKKYSKNSFDAPKNSSKNIFERYIDNYNKWRDGGIDKAIEVIEKERNDVAILSLDLKGFYYNIDIDFDAIDRIIDEEQLEEIKALSMLLNSKL<br>RLMHQKYREKISPYIGVTHSTPSSKGIPIGFTSSAILSNWYLMGFDKDIKSKINPSYYGRYVDDILLVFSSPDIKKEDKGEEVIKFIERTLNGFIKQAANDEKGFTLTESYHELPIQKD<br>KLIFHYFNKDHSLAGLQVFKQEIENRSSAFRFLPEEHISSDLDKFAYDILLDGSANKFRSIVGLAENETELSKYISSHILAHRLCNLPSNENTLKQITLFFKGENSLRFSRLWEKVL<br>SYTLITRKYGFSIAFYKQIKESINKMKWKEGDSVISPKLKNGMQKYIDISLCLNLALLDLNVILGDKEPQNRALEGLKSLIKQSDGMSVLIERFRSSNLIRHNLVSWPLANYTKYKG<br>DLTEENLYSKMHGIPLDNDKLNKTPRYIHADEKQLFDLIDALHARKLHKFTHENKHHLKNCSVTKIKDNKALEIKVGIDDRSGNDTIKVALANMRVERRNIESACRKDQSPNLS<br>YERQRNLYRILNSATEENVQILLLPELSIPVSWLPFMAAHSRRKQISLIFGLEHWVINERAYNILVEMLPYTSNFKHKSSMLVFRVKNHYAPSEITMLNSLRLKISSPKPSKQRYHLI<br>KWKNVSFSTYNCFELANIEHRALFRSQLDILFACVWNQDINYYQHITESATRDLHCYVAQSNTSHYGGSCVLQPTSSTISNKIYVKGGENHCILTTTLNIYALREAQYRSFRINS<br>DTIKHNPPGFDYNALLERGEK |
| DRT5S<br>(UG5S_WP_16609<br>9902.1_Duganella<br>aceris_RC) | MHKPNYQIEDISSAYRKFKSLIYYDKNDLSTRIKLAKFETGLDKKLLNILDVVNYRDPCRHVDFVAWVEDIGSRIVPKGIESEPERTVEENGIFISNFLSYKKLNVKKVNYFFDGPIE<br>LQIIATLWIIHEGKYLDYMLGNECYGSRLESDATLKEESSPKLFIRYHDLYKRWRDGGIKKARQMLVEEKTSVCILGLDIQEYYYHIDFDYKEAKKIIQEKMALESSKNGLIDCLEIIS<br>KKYKSEINSHLPFSHDSDALPTGLPIGMISSPLLANWHLRKFDFEVKEKIRPSYYGRYIDDIFIVIPAPAGFSENKKPIEYFISEILVKNTILKTLPDDRYEINSVPGLYLQQKKCILQH<br>FDINHSIAALEKFKKELEANSSDFLLLPVDDNESSLEDVAYDLLYDGSVNKFRSVKGVAENRYELAKHLARQTMLHLLTTDKFEIDTRNGLLNFFKGKNAIDFHDLWERVLTFFV<br>VSRDAKGFNVFCKNIELEIERINFKSSVDITLLLRENLSTHLNLSKCMASTLDLNADDFDDSRNSLSGKIWTSNLIRHHFVRIPLINYTSFRGSLIGNEASSNLRLNSRKIKYSPRF<br>VNFDECIILITSKILKYHSASSQVDYIVAEAEKLFFRINGRLQSGVEVVKEEEGSDEL |
| AbiP2 | MKKVYELTSEEALSYFLRHDSYTTLELPAYINFTTLLNDINSSIHNKKIKIEPTAKELMGKDINYEVLVSKDGLYSWRRITLINPLYYVYFCRKITAPATWEIITEKFKSFESNDLFTCSSI<br>PVRKDNSSNIAASVMNWWEDFEQKSLALALEYEFMFSTDISNFYPSIYTHSFEWVFISKEEAKKKKSKNNPGGLIDSHIQMMMNNQTNGIPLGSTLMDTFAELILGQIDIELRK<br>KTNELKIINYKVVRYRDDYRIFSNSKDDLDIISKCLVNVLGDFGLDLNSKKTELYEDIILHSLKQAKKDYIKEKRHKSLQKMLYSIYLFSLKHPNSKTTVRYLNDFLRNLFKRKTIKDN<br>GQQVDAMLGIISSIMAKNPTTYPVGTAIFSKLLSFLYGDDTQKKLTKLEQLHKKLDKQPNTEMLDIWFQRTQAKINLEWNKSYKSALCVRINDELTKEKTFSVNNLWNIDWIQGK<br>ETSPNKAKILSLLRKTKIVDTDKFDKMDDNITPEEVNLFFKEHSN |
| T4 Dda | MTFDDLTEGQKNAFNIVMKAIKEKKHHVTINGPAGTGKTTLTKFIIEALISTGGTGIILAAPTHAAKKILSKLSGKEASTIHSILKINPVTYEENVLFEQKEVPDLAKCRVLICDEVSMY<br>DRKLFKILLSTIPPWCTIIGIGDNKQIRPVEPGENTAYISPFFTHKDFYQCELTEVKRSNAPIIDVATDVRNGKWNYDKVVDGHGVRGFTGDTALRDFMVNYFSIVKSLDDLFENR<br>VMAFTNKSVDKLNSIIRKKIFETDKDFIVGEIIVMQEPLFKTYKIDGKPVSEIIFNNGQLVRIIEAEYTSTFVKARGVPGEYLIRHWDLTVETYGDDEYYREKIKIISSDEELYKFNLFLA<br>KTAETYKNWNKGGKAPWSDFWDAKSQFSKVKALPASTFHKAQGMSVDRAFIYTPCIHYADVELAQQLLYVGVTRGRYDVFYV |
| T4 gp32<br>(SSB) | MFKRKSTAELAAQMAKLNGNKGFSSEDKGEWKLKLDNAGNGQAVIRFLPSKNDEQAPFAILVNHGFKKNGKWYIETCSSTHGDYDSCPVCQYISKNDLYNTDNKEYSLVKR<br>KTSYWANILVVKDPAAPENEGKVFKYRFGKKIWDKINAMIAVDVEMGETPVDVTCPWEGANFVLKVKQVSGFSNYDESKFLNQSAIPNIDDESFQKELFEQMVDLSEMTSKD<br>KFKSFEELNTKFGQVMGTAVMGGAAATAAKKADKVADDLDAFNVDDFNTKTEDDFMSSSSGSSSSADDTDLDDLLNDL |

### Extended Data Table 2. Cryo-EM data collection and refinement statistics

| | |
|---|---|
| **Sample** | DRT1 + dNTPs |
| **PDBID** | 9YFD |
| **EMDB code** | EMD-72883 |
| **Microscope** | FEI Titan Krios |
| **Voltage (kV)** | 300 |
| **Detector** | K3 |
| **Magnification (nominal)** | 105,000 |
| **Pixel size (Å/pix)** | 0.826 |
| **Exposure rate ($e^-$/pix/sec)** | 18.7 |
| **Exposure ($e^-$/Å$^2$)** | 55 |
| **Defocus range (mm)** | 0.9–2.4 |
| **Tilt angle (°)** | 0 |
| **Micrographs collected** | 10,188 |
| **Micrographs used** | 3,895 |
| **Particles extracted (total)** | 1,134,805 |
| **Automation software** | SerialEM |
| **3D reconstruction statistics** | |
| **Particles** | 140,634 |
| **Helical parameters** | |
| **Order** | 2 |
| **Rise (Å)** | 59.22 |
| **Twist (°)** | -121.22 |
| **Point group symmetry** | D2 |
| **Map sharpening B-factor** | -88.1 |
| **Unmasked resolution at 0.5 FSC (Å)** | 3.6 |
| **Masked resolution at 0.5 FSC (Å)** | 2.9 |
| **Unmasked resolution at 0.143 FSC (Å)** | 3.1 |
| **Masked resolution at 0.143 FSC (Å)** | 2.6 |
| **Model refinement and validation statistics** | |
| **Composition** | |
| **Non-hydrogen atoms** | 73,480 |
| **Protein residues** | 9,000 |
| **Ligands** | 24 (8 ATP, 16 $Mg^{2+}$) |
| **RMSD bonds (Å)** | 0.002 |
| **RMSD angles (º)** | 0.56 |
| **Average B-factors** | |
| **Protein residues** | 39.4 |
| **Ligands** | 94.0 |
| **Ramachandran** | |
| **Favored (%)** | 97.14 |
| **Allowed (%)** | 2.86 |
| **Outliers (%)** | 0 |
| **Rotamer outliers (%)** | 0.67 |
| **Clash score** | 3.49 |
| **CaBLAM outliers (%)** | 0.58 |
| **CC (mask)** | 0.90 |
| **MolProbity score** | 1.29 |
| **EMRinger score** | 4.45 |

**Extended Data Table 3. nucleotide ion masses used to identify modified peptides[8].**

| Nucleobase | N[H+] | n-$H_2O$ [H+] | dNMP [H+] |
|---|---|---|---|
| **dAdenine** | 136.0618 | 216.0880 | 332.0754 |
| **dGuanine** | 152.0567 | 235.1064 | 348.0704 |
| **dThymine** | 127.0502 | 210.0999 | 323.0639 |
| **dCytosine** | 112.0505 | 195.1002 | 308.0642 |

N: Nucleobase, n: nucleoside, dNMP: deoxyribose nucleotide monophosphate

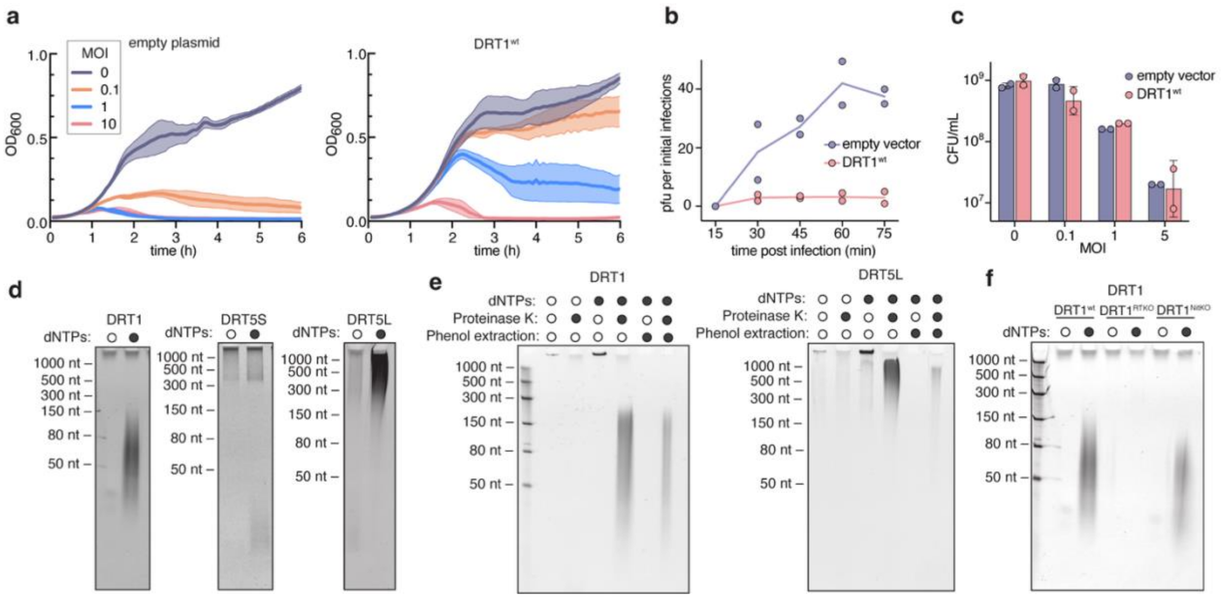


**Extended Data Fig 1. DRT1 antiphage activity and assay of DNA synthesis by purified DRT1, DRT5S, and DRT5L. a)** Growth curve of cells expressing empty plasmid or $DRT1^{wt}$ +/- T4 phage at the indicated multiplicity of infection (MOI). Data are shown as mean ± s.d. for n = 3 independent biological replicates. **b)** Burst size of T4 phage infecting cells with or without $DRT1^{wt}$. Data are shown as mean ± s.d. for n = 2 independent biological replicates per condition. **c)** Cell survival assay of bacteria following 15 min of T4 phage infection. **d)** dNTP polymerization assay of purified DRT1, DRT5S, and DRT5L. Samples were treated with proteinase K, then resolved on 15% TBE-urea gel. **e)** Covalent attachment of DNA to DRT1 and DRT5 as demonstrated by the necessity of proteinase K digestion to release DNA into the soluble phase during phenol-chloroform extraction. **f)** dNTP polymerization assay of purified WT ($DRT1^{wt}$), RT-inactive ($DRT1^{RTKO}$), and nitrilase-inactive ($DRT1^{NitKO}$) protein.

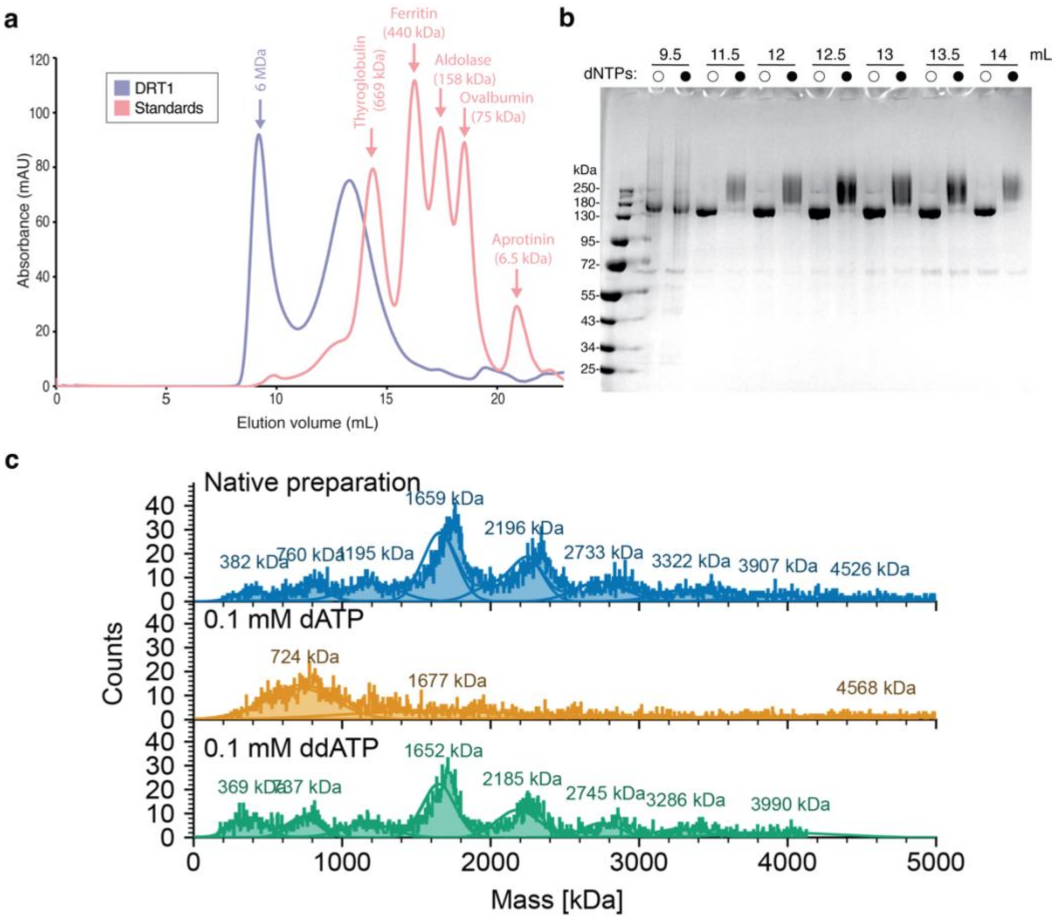


**Extended Data Fig. 2.** Intact, denatured LCMS deconvoluted spectra of **a)** DRT1$^{wt}$, and **b)** DRT1$^{RTKO}$. DRT1$^{wt}$ shows the addition of mass of approximately 312 Da, consistent with deoxyadenosine monophosphate (313 Da) addition. **c)** Reverse phase base peak traces of DRT1$^{wt}$ tryptic digest with extracted ion chromatogram highlighting deoxyadenosine [dA]-containing peptides. Extracted ion chromatograms of the peptide INEIQNEVGTSSIPER (+3 charge state) with 1-4 deoxyadenosines showing sequential elution of the single dA-bound species eluting first, followed by the sequential elution of each +1[dA] species.

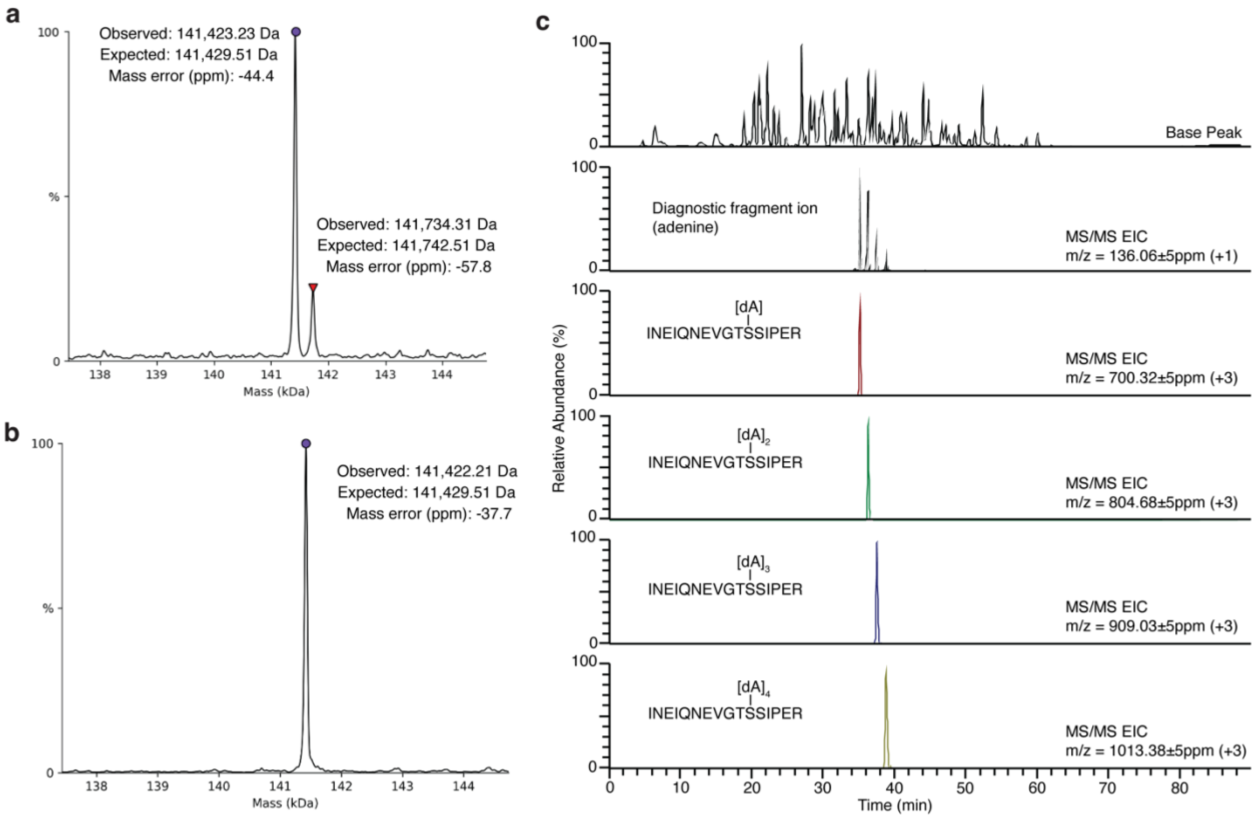


**Extended Data Fig. 3.** HCD product-ion triggered EThcD mass spectrum of DRT1$^{wt}$ tryptic peptide containing a bound deoxyadenosine. (INEIQNEVGTS[dA]SIPER, 2+, m/z 1050.48). A peptide sequence MS/MS coverage map is shown in the upper right corner of the MS/MS spectrum, and a table of observed fragment ions within ±10ppm mass error of the expected mass. Key nucleotide localizing fragments are outlined in purple.

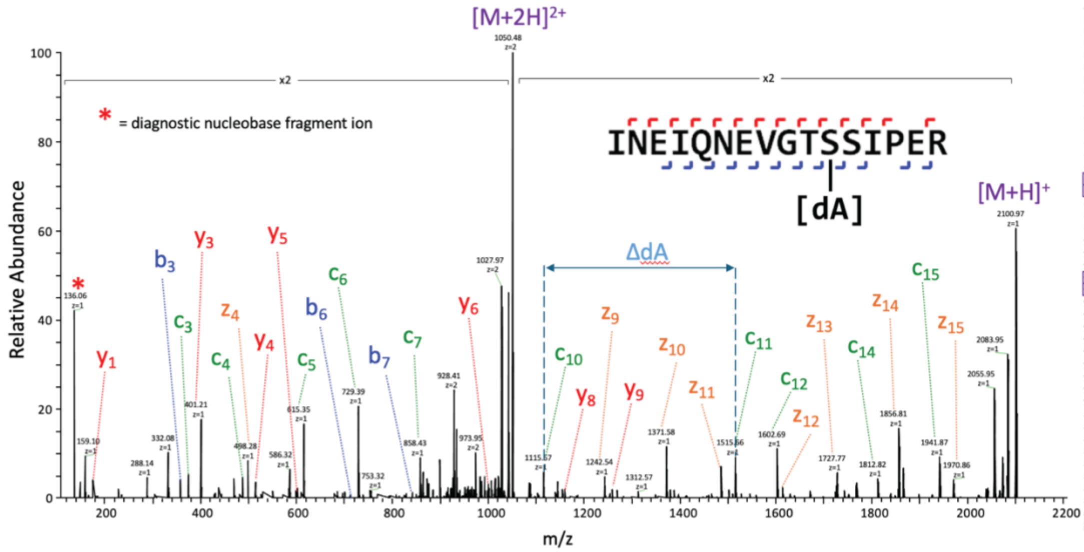


| Fragment Ion | Theoretical Mass | Observed Mass | Mass Error (ppm) |
|---|---|---|---|
| $b_3$ | 356.1696 | 356.1695 | -0.2 |
| $b_5$ | 597.3122 | 597.3127 | 0.8 |
| $b_6$ | 711.3552 | 711.3553 | 0.2 |
| $b_7$ | 840.3977 | 840.3965 | -1.5 |
| $b_8$ | 939.4662 | 939.4645 | -1.8 |
| $c_3$ | 373.1959 | 373.1961 | 0.6 |
| $c_4$ | 486.2799 | 486.2806 | 1.4 |
| $c_5$ | 614.3385 | 614.3390 | 0.8 |
| $c_6$ | 728.3815 | 728.3821 | 0.9 |
| $c_7$ | 857.4240 | 857.4245 | 0.5 |
| $c_8$ | 956.4925 | 956.4908 | -1.7 |
| $c_9$ | 1013.5139 | 1013.5120 | -1.9 |
| $c_{10}$ | 1114.5616 | 1114.5626 | 0.9 |
| $c_{11}$ | 1514.6523 | 1514.6496 | -1.8 |
| $c_{12}$ | 1601.6844 | 1601.6839 | -0.3 |
| $c_{14}$ | 1811.8212 | 1811.8168 | -2.4 |
| $c_{15}$ | 1940.8638 | 1940.8635 | -0.1 |
| $y_1$ | 174.1117 | 174.1114 | -1.6 |
| $y_3$ | 400.2070 | 400.2070 | -0.1 |
| $y_4$ | 513.2911 | 513.2915 | 0.8 |
| $y_5$ | 600.3231 | 600.3238 | 1.1 |
| $y_6$ | 1000.4138 | 1000.4128 | -1.0 |
| $y_7$ | 1101.4615 | 1101.4613 | -0.2 |
| $y_8$ | 1158.4830 | 1158.4822 | -0.7 |
| $y_9$ | 1257.5514 | 1257.5496 | -1.4 |
| $y_{10}$ | 1386.5940 | 1386.5910 | -2.2 |
| $y_{11}$ | 1500.6369 | 1500.6369 | 0.0 |
| $y_{12}$ | 1628.6955 | 1628.6981 | 1.6 |
| $y_{13}$ | 1741.7796 | 1741.7786 | -0.5 |
| $y_{14}$ | 1870.8221 | 1870.8313 | -0.6 |
| $y_{15}$ | 1984.8651 | 1984.8755 | 5.3 |
| $z_4$ | 497.2723 | 497.2728 | 1.0 |
| $z_7$ | 1085.4428 | 1085.4403 | -2.3 |
| $z_8$ | 1142.4642 | 1142.4627 | -1.3 |
| $z_9$ | 1241.5326 | 1241.5310 | -1.3 |
| $z_{10}$ | 1370.5752 | 1370.5748 | -0.3 |
| $z_{11}$ | 1484.6182 | 1484.6173 | -0.6 |
| $z_{12}$ | 1612.6767 | 1612.6739 | -1.8 |
| $z_{13}$ | 1725.7608 | 1725.7588 | -1.2 |
| $z_{14}$ | 1854.8034 | 1854.8026 | -0.4 |
| $z_{15}$ | 1968.8463 | 1968.8364 | -5.0 |

**Extended Data Fig. 4. Oligomeric state of DRT1. a)** Superose 6 10/300 GL size exclusion chromatogram of DRT1$^{wt}$ overlaid with a chromatogram of protein standards and their relative masses. **b)** DRT1$^{wt}$ fractions (elution volume in mL) from **(a)**, either reacted with dNTPs (filled circles) or unreacted (open circles), then resolved by SDS-PAGE. **c)** Mass photometry of DRT1$^{wt}$ native preparation, or reacted with dATP or ddATP.

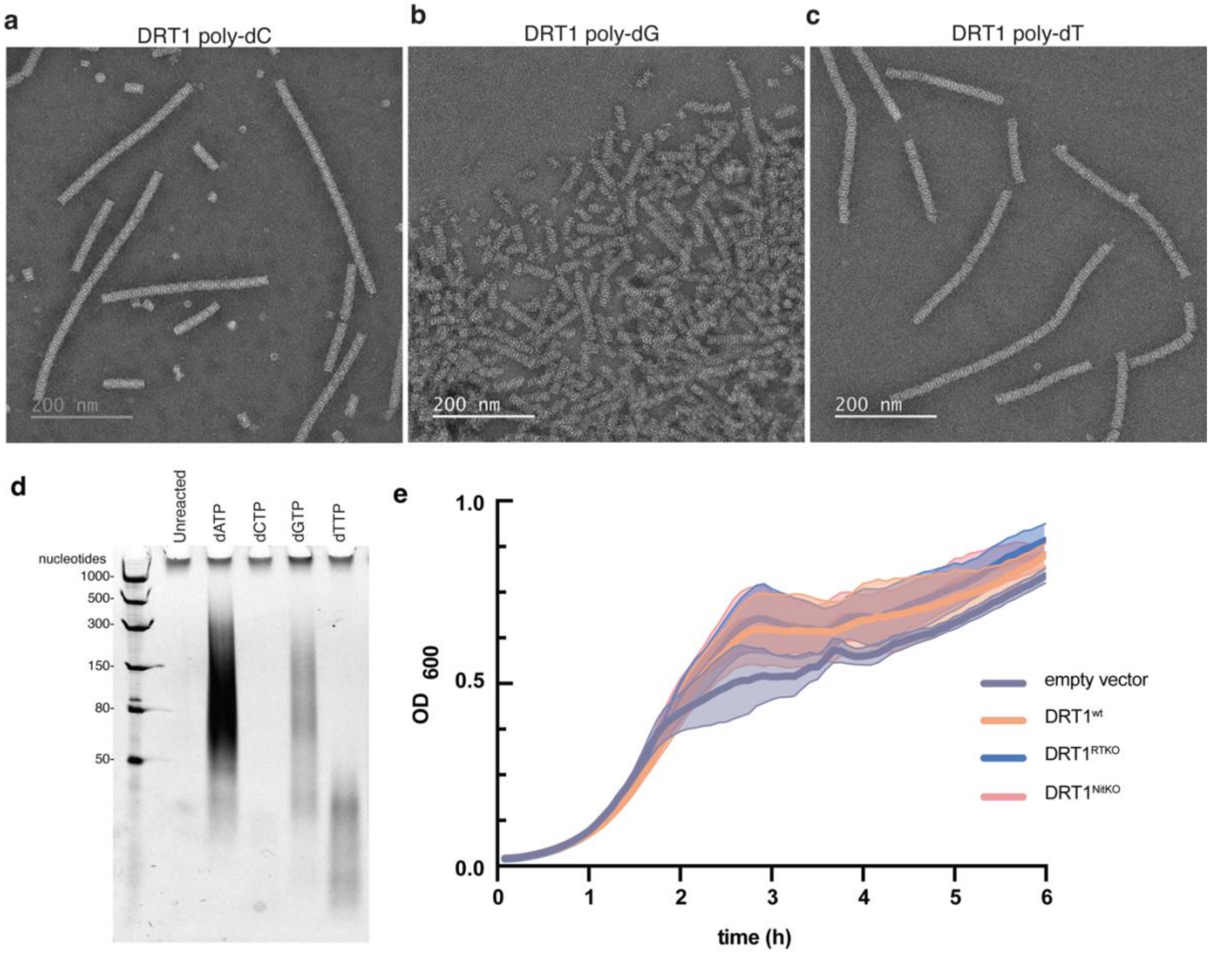


**Extended Data Fig. 5. Filament formation, nucleotide preference, and absence of DRT1 toxicity in the absence of phage. a-c,** Negative stain EM of DRT1$^{wt}$ reacted with **a**) dCTP, **b**) dGTP, or **c**) dTTP. **d)** dNTP polymerization assay gel of DRT1$^{wt}$ reacted with the indicated individual dNTPs. **e)** Growth curve of cells expressing empty plasmid or DRT1 in the absence of phage. Data are shown as mean ± s.d. for n = 3 independent biological replicates.

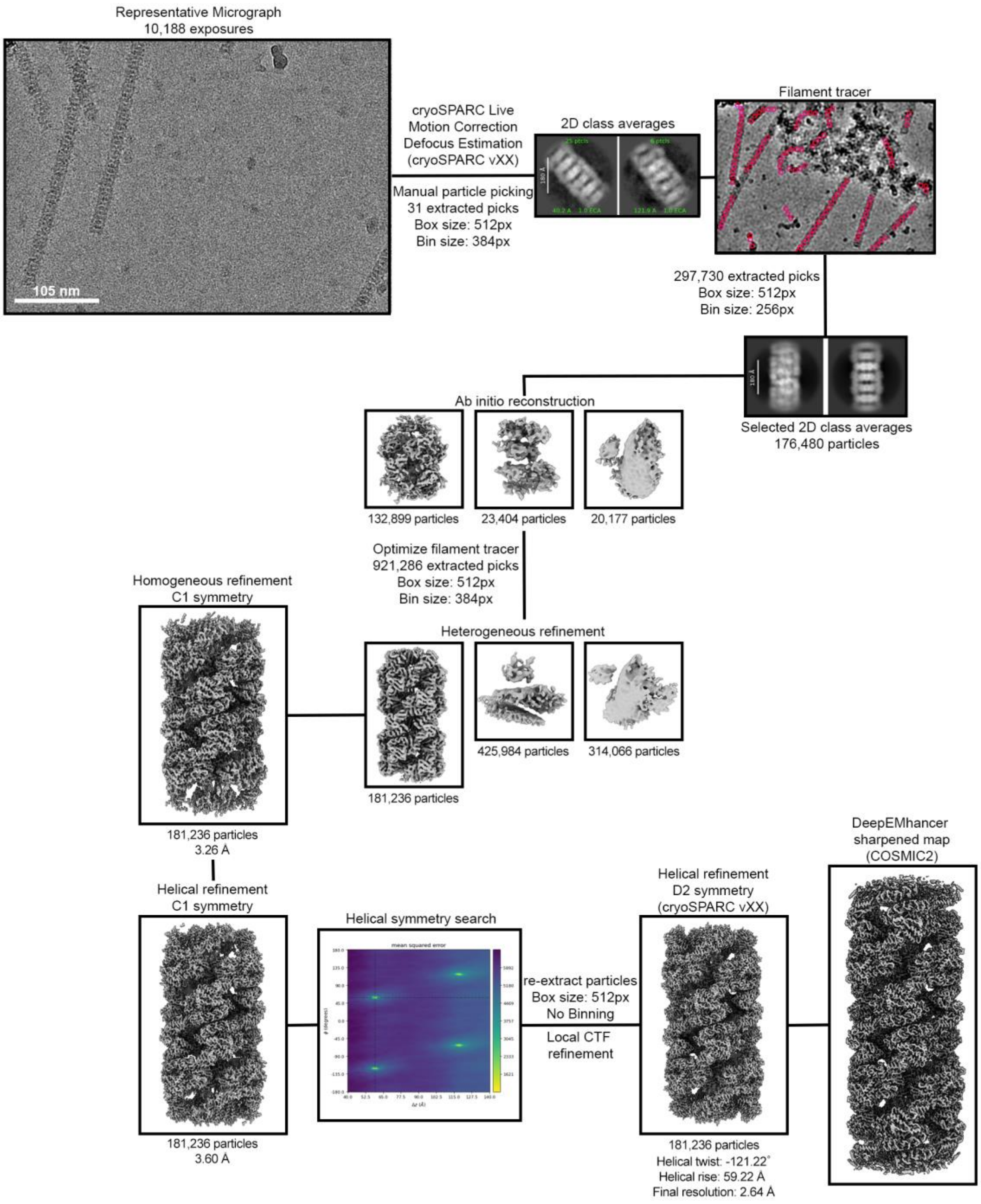


**Extended Data Fig. 6. Cryo-EM data processing workflow.** Flowchart illustrating the cryo-EM data processing steps used to obtain the structure of the DRT1 filament. Particle counts at each stage are indicated, along with resolutions and symmetry parameters for the 3D reconstructions.

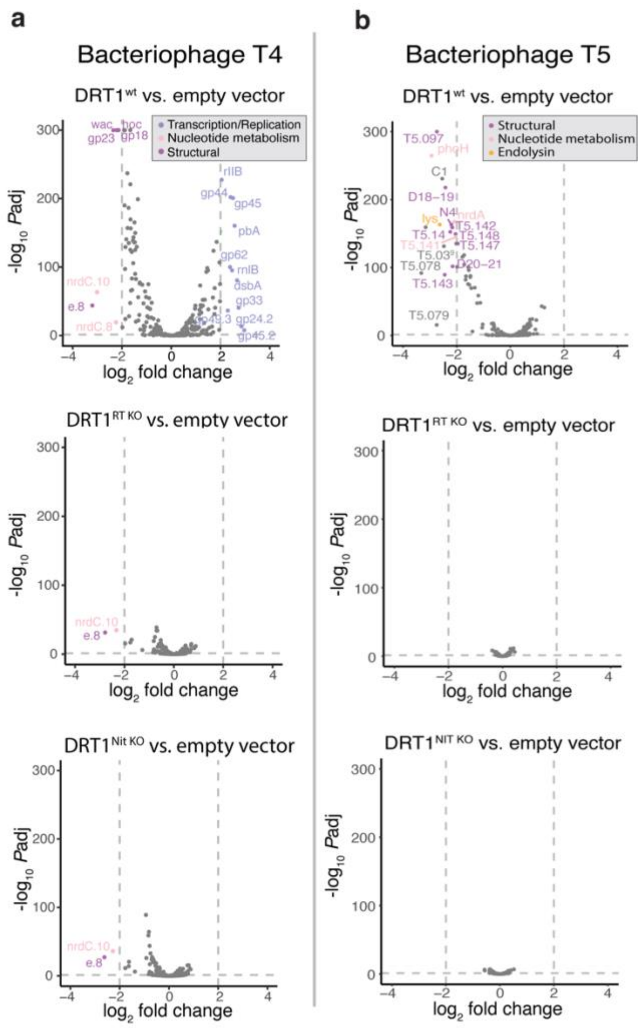


**Extended Data Fig 7. DRT1 alters bacteriophage transcription profiles. a)** Volcano plot showing significantly upregulated and downregulated genes in the bacteriophage T4 genome upon exposure to DRT1$^{wt}$, DRT1$^{RT\ KO}$, DRT1$^{Nit\ KO}$. Genes are colored according to function. **b)** Volcano plot showing significantly upregulated and downregulated genes in the bacteriophage T5 genome upon exposure to DRT1$^{wt}$, DRT1$^{RT\ KO}$, DRT1$^{Nit\ KO}$. Genes are colored according to function.

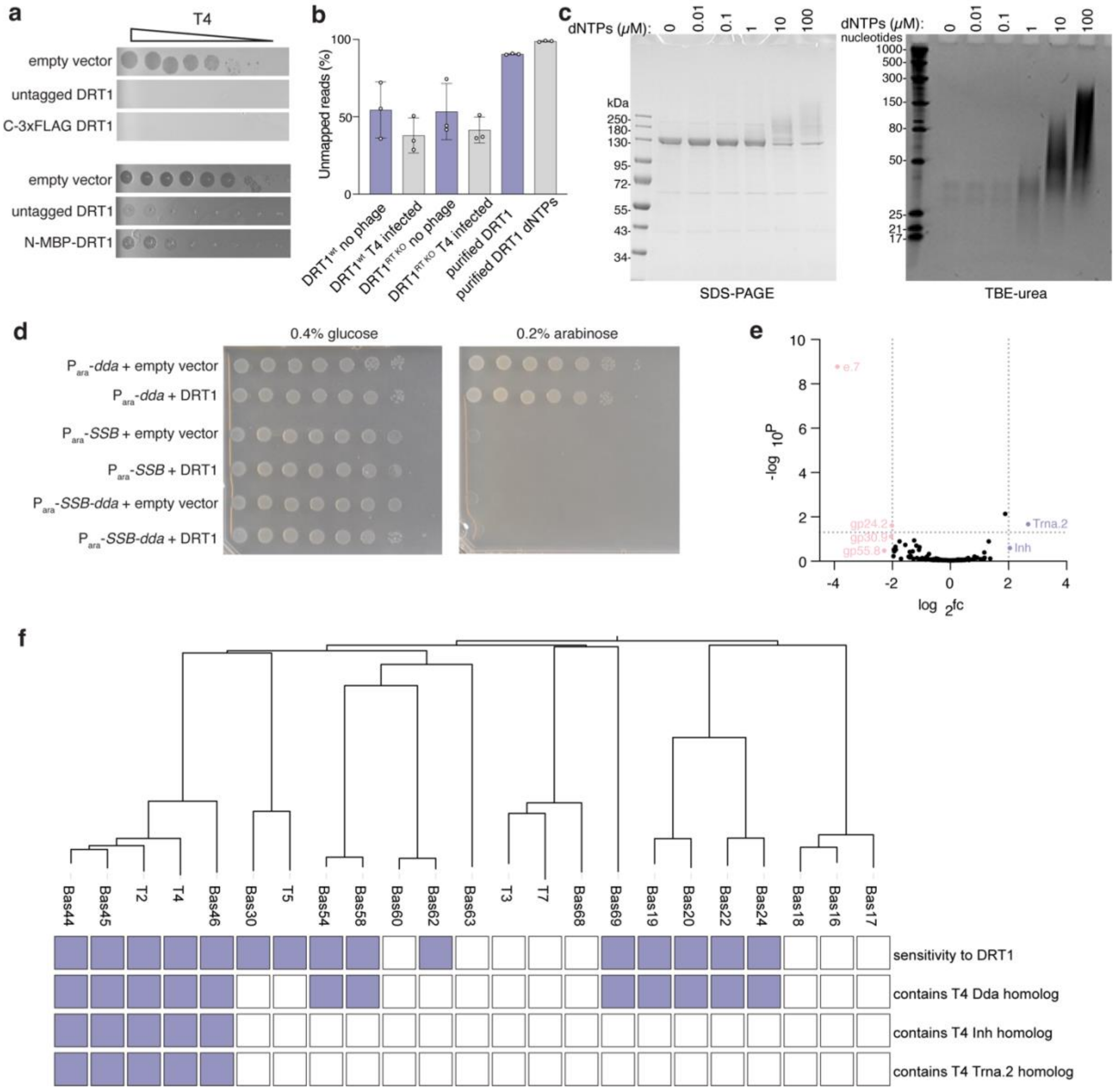


**Extended Data Fig. 8. Viral determinants of DRT1 activation. a)** Phage defence plaque assay of DRT1 with either no tag, a C-terminal 3xFLAG tag, or an N-terminal MBP tag. **b)** Comparison of total unmapped reads between cDIP-seq samples. **c)** Dose-response of DRT1$^{wt}$ to dNTP mix titration. Samples run directly on SDS-PAGE (left) or digested with proteinase K and resolved on TBE-urea (right). **d)** Co-toxicity cell viability assay of DRT1$^{wt}$ with candidate T4 phage trigger genes expressed from an arabinose-inducible promoter ($P_{ara}$). **e)** LC-MS/MS proteomic analysis, volcano plot of differentially enriched T4 phage proteins following infection of cells expressing DRT1$^{wt}$ versus DRT1$^{RT\ KO}$. **f)** Phylogenetic tree of phage collection used in this study with indicators of sensitivity to DRT1, presence of Dda, Inh, Trna.2 or homologs.
